# A Multisensor Framework Reveals Redox Constraints on Glycolysis *in vivo*

**DOI:** 10.64898/2026.06.16.732715

**Authors:** Snusha Ravikumar, Aaron D. Wolfe, Daniel A. Colón-Ramos

## Abstract

Genetically encoded biosensors have transformed the study of metabolism, yet measurements of individual metabolites often provide an incomplete view of pathway regulation. Here, we develop a multisensor framework in *Caenorhabditis elegans* neurons to interpret glycolytic dynamics and redox state *in vivo*. We combine biosensors for NADH/NAD⁺, fructose-1,6-bisphosphate, lactate, and pyruvate to resolve metabolic responses during hypoxia and redox perturbation.

To causally test how redox state modulates glycolysis *in vivo*, we cell-specifically expressed the NADH-producing enzyme *Ec*STH and the NADH oxidase *Lb*NOX to bidirectionally tune neuronal NADH/NAD⁺ balance. These perturbations revealed that redox modulation is sufficient to constrain or relieve lower glycolytic activity. Elevation of NADH/NAD⁺ promoted accumulation of upper glycolytic intermediates while suppressing lower glycolytic responses during energetic stress, consistent with inhibition at the NAD⁺-dependent GAPDH step. Conversely, oxidation of NADH relieved this constraint and shifted metabolite pools consistent with enhanced lower glycolytic activity. Elevated NADH/NAD⁺ ratios also impaired synaptic vesicle organization, linking redox-mediated glycolytic inhibition to neuronal function.

As a case study for how integrated biosensor approaches can provide semi-quantitative insight into pathway-level metabolic regulation, we genetically perturbed endogenous NADH recycling pathways. These experiments revealed a hierarchical organization of neuronal redox buffering, with lactate dehydrogenase (LDH-1) serving as the dominant route for NAD⁺ regeneration during hypoxia and glycerol-3-phosphate dehydrogenase (GPDH-2) providing a secondary compensatory pathway. Graded impairment of NADH recycling resulted in corresponding increases in fructose-1,6-bisphosphate accumulation and synaptic defects, consistent with progressive inhibition of lower glycolysis. Together, these results establish a tractable *in vivo* system to probe causal relationships between redox state, glycolytic dynamics, and cellular physiology.

## INTRODUCTION

Neuronal activity is energetically expensive. Maintenance of ion gradients, synaptic vesicle recycling, and neurotransmission impose continuous ATP demand that must be dynamically matched by metabolic supply^1–3^. While individual metabolic pathways have been extensively characterized biochemically, understanding how energetic state enables or constrains neuronal function within an intact nervous system remains a central challenge. It is increasingly clear that neurons must rely on distinct, cell-autonomous glycolytic capacities to meet localized energetic demands^4–6^. *In vivo*, neurons must coordinate ATP production, redox balance, and biosynthetic flux across diverse cell types and subcellular compartments, while simultaneously maintaining circuit-level stability. Linking energetic state to neuronal physiology therefore requires resolving metabolic dynamics at cellular and subcellular resolution *in vivo*.

Despite major advances in metabolic imaging, the spatiotemporal dynamics of metabolism remain incompletely resolved. Approaches such as functional magnetic resonance imaging (fMRI)^7^ and imaging mass spectrometry^8–10^ provide valuable systems-level insights but lack the cellular or temporal resolution required to dissect pathway-specific flux within individual neurons undergoing physiological changes. Shifts in cytosolic redox state (NADH/NAD+ ratios), initially detected through intrinsic NADH fluorescence and now measured by FLIM^11,12^ or genetically encoded sensors^13,14^ are frequently used to indirectly infer glycolytic activity^4^. More recently, biosensors such as HYlight have enabled direct measurement of the upper glycolytic intermediate fructose-1,6-bisphosphate (FBP)^5,15^. In contrast to bulk biochemical approaches, genetically encoded biosensors enable metabolic measurements at cellular and subcellular resolution *in vivo* and have revealed metabolic heterogeneity and dynamics that would otherwise remain obscured^4,5^. Sensors for metabolites associated with lower glycolysis have also been developed^16–19^, such that measurements now span upper glycolysis, lower glycolysis, and cytosolic redox state. Despite this expanded toolkit, interpretation of biosensor dynamics remains challenging. Biosensors are often applied individually and in distinct experimental contexts, making it difficult to infer pathway-level behavior from any single readout alone. An increase in a metabolite can reflect multiple underlying metabolic states, including increased production, reduced downstream consumption, or altered metabolite exchange between pathways or cells. As a result, single-metabolite measurements can be insufficient to resolve glycolytic state *in vivo*.

In addition, interpretation of redox measurements presents a unique challenge. An elevated NADH/NAD⁺ ratio can be interpreted as a signature of active glycolysis^4^, yet in specific conditions, it can simultaneously impose a biochemical limitation on the pathway^20–23^. The lower glycolytic enzyme glyceraldehyde-3-phosphate dehydrogenase (GAPDH) requires NAD⁺ as an obligate cofactor^24,25^. Thus, glycolytic flux both generates NADH and depends on continuous NAD⁺ regeneration. Without efficient NADH recycling, relative NAD⁺ depletion restricts GAPDH and stalls lower glycolysis. Consequently, measurements of redox state or upstream intermediates alone cannot distinguish whether glycolysis is accelerating or metabolically bottlenecked by cofactor imbalance. Resolving glycolytic function therefore requires integrating measurements across multiple steps of the pathway.

Finally, establishing causal relationships between glycolysis and cellular function requires tools that move beyond metabolic observation and enable targeted metabolic manipulation. Coupled with live-cell imaging, such perturbations allow direct assessment of how changes in glycolysis impact cellular physiology. Among glycolytic regulatory nodes, cytosolic redox state represents a potential lever: because the NAD⁺-dependent GAPDH reaction links lower glycolysis directly to NADH/NAD+ state, shifts in the NADH/NAD⁺ ratio may gate glycolytic flux without altering glycolytic enzymes themselves. This coupling raises a testable hypothesis: that targeted manipulation of neuronal NADH/NAD⁺ ratios is sufficient to modulate glycolytic flux in a cell-autonomous manner^26^. Genetically encoded redox modulators provide a strategy to directly test this idea^27^. *Lb*NOX lowers the NADH/NAD⁺ ratio by oxidizing NADH, whereas *Ec*STH increases NADH through transhydrogenase activity^28,29^. Although these tools have been used to probe redox biology in cultured cells and select *in vivo* systems^30–32^, they have not been used to quantitatively tune glycolytic flux *in vivo*.

Here, we address these challenges by developing an *in vivo* multisensor framework to dissect the redox architecture of neuronal glycolysis in *Caenorhabditis elegans*. By sequentially integrating genetically encoded reporters of upper glycolysis, lower glycolysis, and cytosolic redox state within single neurons, we establish a pathway-level view of glycolytic dynamics that cannot be obtained from individual metabolite measurements alone. We further combine these measurements with cell-specific genetic perturbations of NADH/NAD⁺ balance to directly test how redox state shapes glycolytic function. Using this approach, we show that elevation of the NADH/NAD⁺ ratio selectively restricts lower glycolysis, whereas enhanced NADH oxidation relieves this constraint, revealing cytosolic redox balance as both a readout and regulator of neuronal glycolysis.

Using this framework, we genetically perturbed endogenous NADH recycling pathways, including lactate dehydrogenase and glycerol-3-phosphate dehydrogenase. These experiments revealed a hierarchical organization of neuronal redox buffering, with lactate dehydrogenase functioning as the dominant source of NAD⁺ regeneration and glycerol-3-phosphate dehydrogenase providing a secondary compensatory pathway. Across perturbations, the extent of FBP accumulation closely tracked NADH/NAD⁺ elevation and associated synaptic defects, demonstrating that cytosolic redox state quantitatively tunes glycolytic dynamics *in vivo*.

## RESULTS

### Live imaging of Lactate dynamics in C. elegans neurons

To interpret glycolytic dynamics in neurons *in vivo*, we sought to integrate multiple genetically encoded biosensors into a framework to monitor distinct stages of glycolysis within the same cellular context **(Figure 1A**). We previously adapted the biosensor HYlight for use in *C. elegans* to monitor fructose-1,6-bisphosphate (FBP) levels, the product of the rate-limiting phosphofructokinase step catalyzed by PFK-1.1 **(Figure 1A)**^5^. However, because accumulation of any single metabolite can reflect either increased flux or impaired downstream consumption, interpretation of FBP changes alone can be ambiguous. Rather than multiplexing multiple biosensors simultaneously within the same animal, which remains technically challenging due to overlapping spectral requirements, we sequentially assayed complementary biosensors in the same identified neuron under the same metabolic perturbation. This strategy enabled comparison of upper glycolytic activity, downstream glycolytic state, and cytosolic redox balance within a common physiological context.

**Figure 1.**
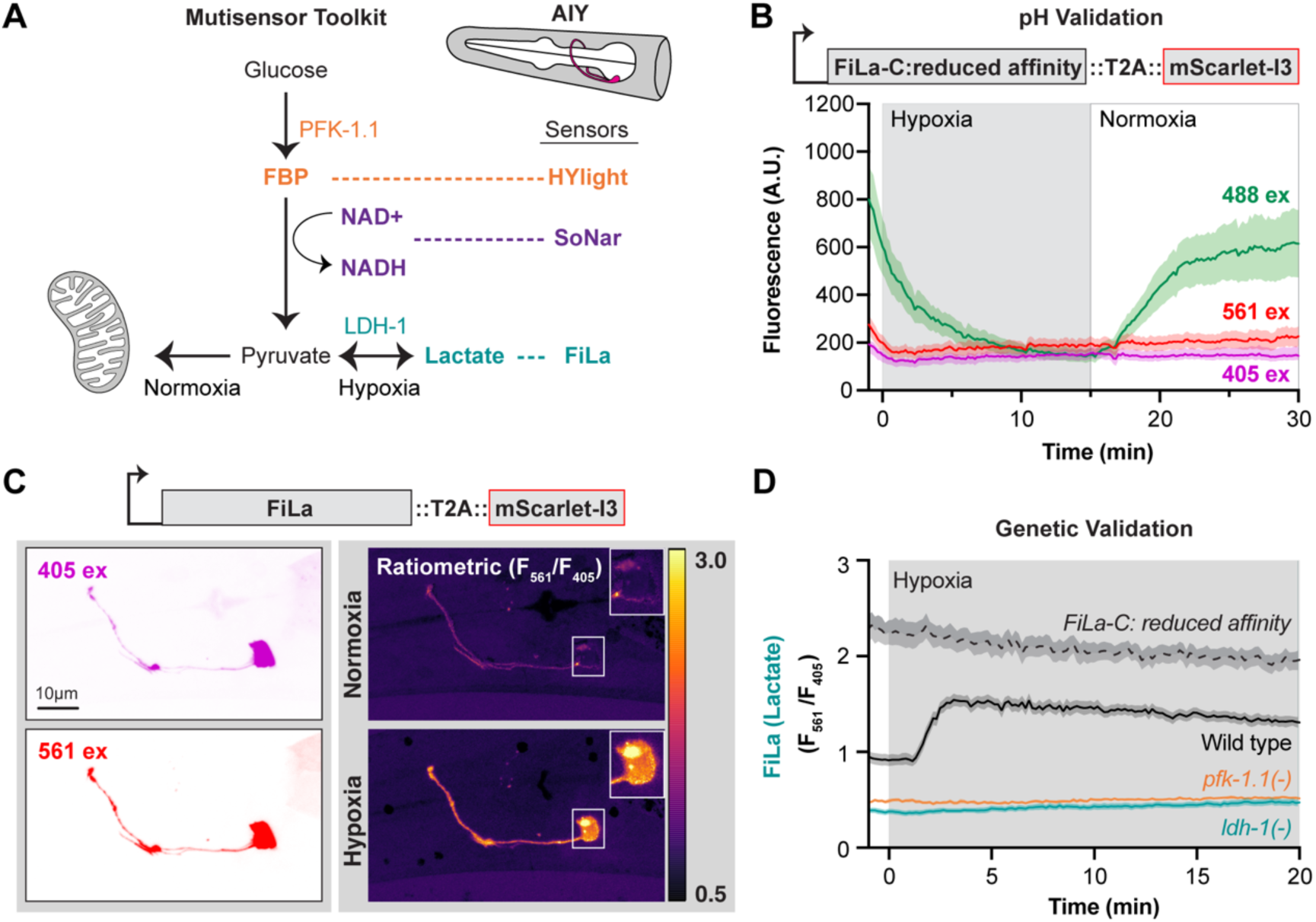
Live imaging of lactate dynamics in *C. elegans* neurons. **(A)** Schematic of glycolysis and associated biosensors used to monitor distinct metabolites in neurons *in vivo*. HYlight reports the upper glycolytic intermediate fructose-1,6-bisphosphate (FBP), SoNar reports cytosolic NADH/NAD⁺ ratios, and FiLa reports lactate levels downstream of pyruvate. Lactate production through LDH-1 is favored under hypoxic conditions, whereas pyruvate can be directed toward mitochondrial metabolism under normoxia. **(B)** Fluorescence intensity of the reduced-affinity lactate sensor FiLa-C under 488 nm (green), 405 nm (purple), and 561 nm (red) excitation during hypoxia and reoxygenation. FiLa-C exhibited strong signal changes under 488 nm excitation but minimal changes under 405 nm and 561 nm excitation. Shaded regions represent mean ± SEM. **(C)** Expression of the lactate biosensor FiLa::T2A::mScarlet-I3 in AIY neurons. Fluorescent images were acquired under 405 nm excitation (*Top Left*) and 561 nm excitation (*Bottom Left*). Ratiometric images (F561/F405) were acquired under normoxia (*Top Right*) and hypoxia (*Bottom Right*). Color scale indicates relative lactate levels. Scale bar, 10 μm. **(D)** Time course of lactate dynamics in AIY neurons measured using FiLa (F561/F405) during 20 minutes of hypoxia. Wild-type animals (black) show an increase in signal upon hypoxia. *pfk-1.1(−)* (orange) and *ldh-1(−)* (teal) mutants exhibit reduced baseline signal and no hypoxia-induced increase. The reduced-affinity control FiLa-C (dashed) shows elevated baseline signal without stimulus-dependent changes. Shaded regions represent mean ± SEM.

Pyruvate generated downstream of FBP is reversibly converted to lactate by lactate dehydrogenase (LDH-1 in *C. elegans*), establishing a near-equilibrium reaction that regenerates NAD⁺ and couples glycolytic flux to cytosolic redox state **(Figure 1A).** We therefore used lactate dynamics alongside FBP measurements as a proxy for glycolytic flux. To monitor lactate dynamics *in vivo*, we evaluated two genetically encoded lactate biosensors with distinct binding affinities and fluorescence properties: FiLa^17^, a high-affinity cpYFP-based sensor, and R-iLACCO1.2^18^, a lower-affinity cpmApple-based sensor **(Supplementary Figure 1F)**. Because FiLa exhibited robust dynamic responses within the physiological lactate range of AIY neurons, we focused initial analyses on this sensor. The circularly permuted YFP (cpYFP)-based sensor was codon-optimized, engineered with a synthetic intron, and expressed in AIY neurons under the *ttx-3* promoter. We validated the sensor in three steps: first by characterizing pH-dependent responses during transient hypoxia, which is known to inhibit mitochondrial OXPHOS and increase glycolytic flux; second by establishing a ratiometric imaging strategy that compares changes in the sensors to a background cytosolic signal; and third by genetically validating lactate-dependent signal changes via CRISPR-engineered mutations of LDH-1.

Similar to other cpYFP based sensors, FiLa’s fluorescence upon excitation at 488 nm has been reported to be sensitive to intracellular pH. Hence it was important to first establish the magnitude of pH changes that occur during our transient hypoxia paradigm. To address this, we leveraged a microfluidic system that enables rapid and reversible exchange between normoxic and hypoxic conditions by flowing oxygen or nitrogen over immobilized animals. To benchmark pH dynamics, we expressed a genetically encoded pH sensor, pHmScarlet, in AIY neurons and measured fluorescence responses during normoxia, hypoxia, and reoxygenation **(Supplementary Figure 1A)**^33^. We observed pronounced fluorescence changes during hypoxia and reoxygenation, consistent with transient cytosolic acidification and recovery. Consistent with lactate production contributing to hypoxia-induced acidification, pH sensor responses were markedly attenuated in *ldh-1(−)* mutants, exhibiting only modest residual fluctuations during hypoxia and reoxygenation **(Supplementary Figure 1A).** These results indicate that transient hypoxia induces a lactate-dependent decrease in cytosolic pH in AIY neurons. Because pH can influence the fluorescence of cpYFP-based biosensors, these changes represent a potential confounding factor when interpreting FiLa responses during hypoxia. To assess the contribution of pH to the FiLa signal, we next leveraged a reduced-affinity FiLa variant (FiLa-C). We reasoned that changes in FiLa-C due to transient hypoxia would represent changes due to extraneous factors, such as pH, and not true changes in lactate levels. FiLa and FiLa-C support ratiometric imaging via excitation at 405 nm and 488 nm. Upon hypoxia, FiLa-C exhibited pronounced fluorescence changes at 488 nm that closely resembled responses measured with the pH sensor **(Figure 1B)**. In contrast, excitation of FiLa-C at 405 nm remained stable across conditions, consistent with prior reports^17^. These results indicate that the 488 nm excitation signal is highly sensitive to the pH changes that accompany hypoxia, whereas the 405 nm excitation signal remains largely pH insensitive under our experimental conditions. Given the substantial lactate-dependent acidification observed during hypoxia, we therefore used 405 nm excitation for all subsequent FiLa measurements.

However, reliance on a single excitation wavelength precluded conventional ratiometric imaging, which is typically used to correct for differences in biosensor expression levels between animals. To address this limitation, we fused FiLa and FiLa-C to the fluorophore mScarlet-I3 using a T2A self-cleaving peptide **(Figure 1B)**. This design permitted computation of the F405/F561 ratio, which in the reduced affinity control FiLa-C remained flat across normoxia and hypoxia, indicating the absence of both stimulus-dependent and imaging-related fluctuations **(Supplementary Figure 1B)**. To account for differing photobleaching rates between the sensor and mScarlet-I3, we quantified fluorescence decay under unstimulated conditions **(Supplementary Figure S2A)**. Fluorescence excited at 405 nm exhibited negligible photobleaching during 20 min of imaging, whereas the 561 nm channel decreased by approximately 11% (slope = 0.006 min⁻¹; **Supplementary Figure S2A**). Therefore, bleaching correction was applied only to the 561 nm channel for FiLa and FiLa-C.

Lactate binding to FiLa reduces fluorescence upon 405 nm excitation; accordingly, lactate accumulation results in a decrease in F405 signal. To facilitate interpretation of the lactate dynamics, we report it as the inverse ratio (F561/F405), such that increases in the reported signal correspond to increases in intracellular lactate. Representative fluorescence images acquired at 405 nm and 561 nm excitation, together with corresponding ratiometric images during normoxia and hypoxia, demonstrated robust lactate-dependent signal changes in AIY neurons, consistent with enhanced anaerobic glycolysis upon hypoxia **(Figure 1C)**. To genetically disrupt lactate production, we generated full-gene deletion alleles, *ldh-1(ola514)* and *pfk-1.1(ola458)*, using CRISPR/Cas9 gene editing. Both mutations significantly reduced the baseline FiLa signal and abolished hypoxia-induced lactate accumulation **(Figure 1D)**. As previously reported, FiLa-C displayed elevated baseline fluorescence^17^ but failed to show lactate induced responses upon hypoxia **(Figure 1D)**.

To determine whether lactate levels vary across neuronal types, we next examined the FiLa signal in a bilaterally asymmetrical sensory neuron pair, ASEL and ASER (**Supplementary Figure S1C**). In contrast to AIY, both ASEL and ASER exhibited substantially elevated steady-state lactate levels **(Supplementary Figure S1D)**. This elevated baseline was abolished in *ldh-1(-)* mutants, consistent with the elevated baseline signal representing differences in endogenous lactate levels across cell types. Surprisingly, upon hypoxia, the FiLa signal in ASEL and ASER did not increase **(Supplementary Figure S1E)**. Because of the high levels of baseline lactate in the ASER and ASEL neurons, we reasoned that the lack of response to transient hypoxia when using the FiLa sensor could be due to a saturation of the dynamic range under basal conditions. We therefore adapted a second lactate biosensor, R-iLACCO1.2^18^, which has a lower affinity for lactate (Kd ∼4 mM) compared to FiLa (Kd ∼130 μM) **(Supplementary Figure S1F)**. Although R-iLACCO1.2 is reported to exhibit pH sensitivity, intracellular acidification during hypoxia would be expected to decrease fluorescence, while increases in lactate would be expected to increase fluorescence, allowing for interpretation of changes in signal dynamics.

When expressed in ASEL and ASER, R-iLACCO1.2 revealed differences in baseline lactate levels between the two neurons. These differences were abolished in *ldh-1(-)* mutants **(Supplementary Figure S1G, *Left*)**. Use of the lower affinity R-iLACCO1.2 sensor also resulted in reproducible hypoxia-induced increases in sensor signal in both neurons **(Supplementary Figure S1G, *Right*)**, consistent with the idea that FiLa was operating near saturation in ASER and ASEL. Together, the optimization of these sensors for use in single neurons and *in vivo* underscores the importance of accounting for baseline metabolite concentrations, sensor affinity, pH sensitivity, and photobleaching, as well as validating sensor responses through genetic perturbations. Moreover, our findings reveal that steady-state lactate levels vary substantially across neuron subtypes, consistent with different metabolic programming across cell types.

### Live imaging of NADH/NAD+ in C. elegans neurons

To visualize neuronal redox dynamics *in vivo*, we adapted the NADH/NAD⁺ biosensor SoNar for use in *C. elegans*^14^ **(Figure 2A)**. Similar to our strategy with FiLa, SoNar was fused to mScarlet-i3 using a T2A self-cleaving peptide to enable ratiometric imaging and correction for differences in expression level, z-drift and animal movement. SoNar is a cpYFP-based biosensor that supports excitation at both 405 nm and 488 nm. As with other cpYFP-based sensors, fluorescence excited at 488 nm has been reported to be sensitive to pH, whereas excitation at 405 nm is comparatively pH-resistant^14^.

**Figure 2.**
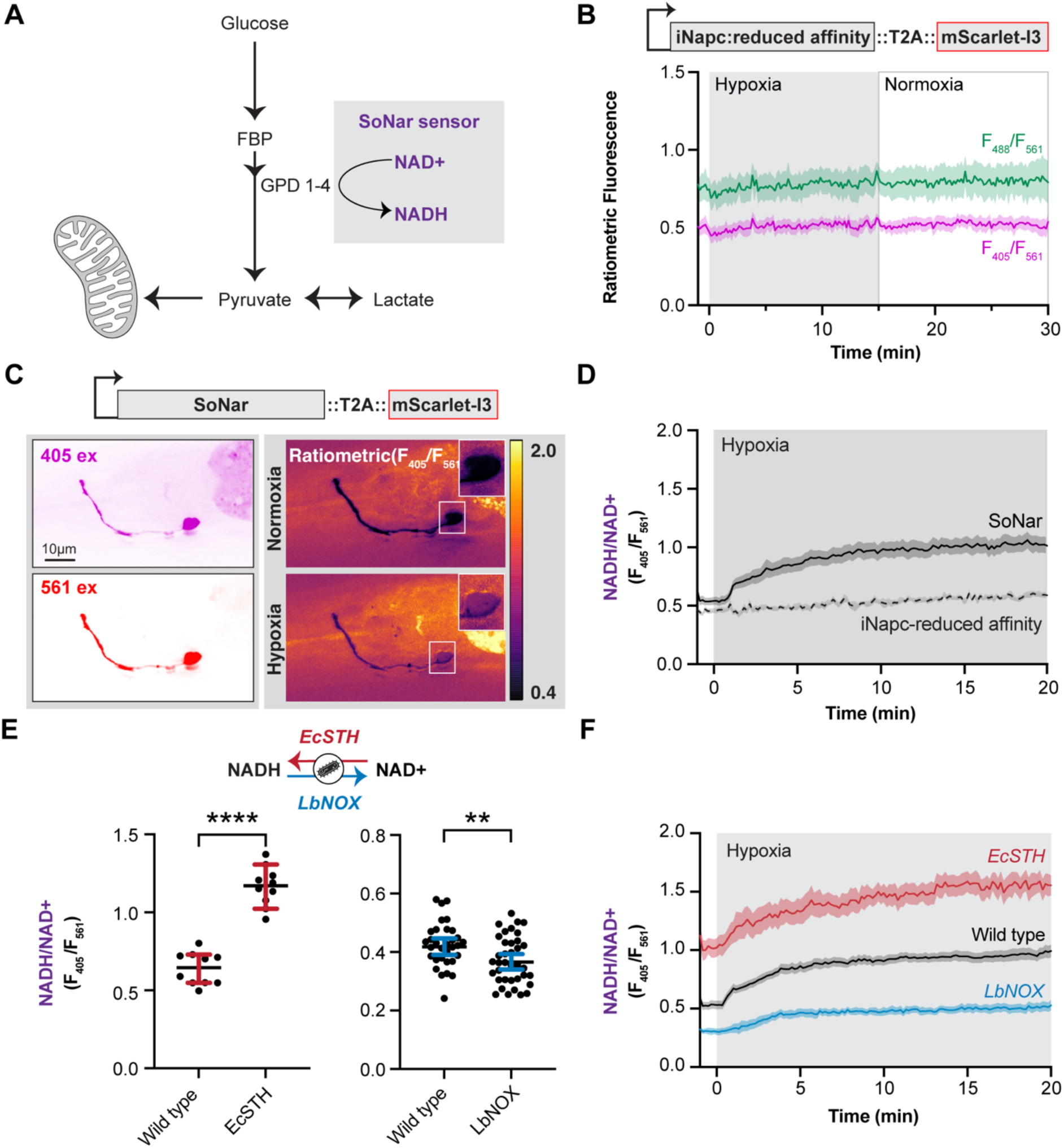
Live imaging of NADH/NAD⁺ ratios in *C. elegans* neurons. **(A)** Schematic of glycolysis highlighting cytosolic NADH/NAD⁺ generation at the GAPDH step, monitored using the biosensor SoNar. GAPDH homologs in *C. elegans* include *gpd-1*, *gpd-2*, *gpd-3*, and *gpd-4*. **(B)** Ratiometric signal of the reduced-affinity NADH/NAD⁺ sensor iNapc calculated as F488/F561 (green) and F405/F561 (purple) during hypoxia and reoxygenation. Shaded regions represent mean ± SEM **(C)** Expression of the NADH/NAD⁺ biosensor SoNar::T2A::mScarlet-I3 in AIY neurons. Fluorescent images were acquired under 405 nm excitation (*Top Left*) and 561 nm excitation (*Bottom Left*). Ratiometric images (F405/F561) are shown under normoxia (Top Right) and hypoxia (*Bottom Right*). Scale bar, 10 μm. **(D)** Time course of NADH/NAD⁺ dynamics in AIY neurons measured using SoNar (F405/F561) during 20 minutes of hypoxia. Wild-type animals (black) exhibit an increase in signal upon hypoxia. The reduced-affinity control iNapc (dashed) shows no stimulus-dependent change. Shaded regions represent mean ± SEM. **(E)** Quantification of steady-state NADH/NAD⁺ ratios (F405/F561) in AIY neurons expressing *Ec*STH (red, *Left*) or *Lb*NOX (blue, *Right*) compared to wild type (black). *Ec*STH expression increases NADH/NAD⁺ ratios, whereas *Lb*NOX expression decreases NADH/NAD⁺ ratios. Each dot represents a single animal. Bars represent mean ± 95% CI. **(F)** Time course of NADH/NAD⁺ dynamics (F405/F561) in AIY neurons during hypoxia in wild-type (black), *Ec*STH-expressing (red), and *Lb*NOX-expressing (blue) animals. Shaded regions represent mean ± SEM. Statistical comparisons were performed using an unpaired Student’s t test. **** p < 0.0001; ** p < 0.01.

To assess sensor specificity and potential pH contributions, we examined iNapc, a SoNar-derived reduced-affinity variant with strongly diminished NADH/NAD⁺ binding^34^. iNapc showed no detectable changes between normoxic and hypoxic conditions at either excitation wavelength, remaining flat at both 405 nm and 488 nm **(Figure 2B),** suggesting that iNapc fluorescence, and therefore SoNar, does not change significantly upon pH changes. Still, we conservatively treated 488-nm excitation of SoNar as potentially pH-sensitive and based quantitative analyses on the 405 nm excitation channel. NADH/NAD⁺ dynamics were quantified using the F405/F561 ratio, in which the 405 nm excitation channel was normalized to the mScarlet-i3 reference 561 nm channel. To account for differing photobleaching rates between the sensor and mScarlet-I3, we quantified fluorescence decay during 20 min of continuous imaging under normoxic conditions **(Supplementary Figure S2B)**. Fluorescence excited at 405 nm and 561 nm decreased by approximately 6% and 14%, respectively, over the imaging period (slopes = 0.003 min⁻¹ and 0.007 min⁻¹). Accordingly, bleaching correction based on the measured decay profiles was applied to both channels for iNapc and SoNar prior to calculation of ratiometric signals. Using the F405/F561 ratio, we detected increases in NADH/NAD⁺ during hypoxia, confirming SoNar responsiveness to acute metabolic stress **(Figure 2C,D)**. The absence of hypoxia-induced responses in iNapc confirmed that SoNar signals reflect changes in NADH/NAD⁺ **(Figure 2D)**.

We next tested whether SoNar could detect controlled bidirectional shifts in cytosolic NADH/NAD⁺ balance. To directly manipulate neuronal redox state *in vivo*, we heterologously expressed two bacterial enzymes in AIY neurons: *E. coli* soluble transhydrogenase (*Ec*STH)^28^, which drives NADH production, increased SoNar signals, whereas *Lactobacillus brevis* NADH oxidase (*Lb*NOX)^29^, which oxidizes NADH, reduced them **(Figure 2C,D)**. These manipulations preserved the hypoxia-induced rise in NADH/NAD⁺. Together, these results establish a pH-insensitive, ratiometric version of SoNar for *C. elegans* neurons and validate its capacity to detect redox changes driven by metabolic stress or targeted enzymatic modulation.

### Elevated NADH/NAD⁺ ratios constrain glycolytic dynamics and disrupt synaptic vesicle organization

Previous work established *Ec*STH as a tool to elevate cytosolic NADH/NAD⁺ ratios and induce reductive stress, leading to increased lactate/pyruvate ratios and accumulation of upper glycolytic intermediates in cultured cells^28^. However, how elevated NADH/NAD⁺ ratios dynamically reshape glycolytic state *in vivo*, and whether these metabolic changes are sufficient to impair physiological outputs, remains unclear. We expressed *Ec*STH in neurons and monitored FBP and lactate levels.

Under baseline normoxic conditions, *Ec*STH expression increased FBP levels relative to wild-type animals **(Figure 3A)**, indicating accumulation of upstream glycolytic intermediates. These findings suggest that elevated NADH/NAD⁺ is sufficient to alter glycolytic state even in the presence of oxygen, although the underlying mechanism cannot be inferred from FBP measurements alone. To examine pathway dynamics during further energetic stress, we next quantified responses to hypoxia. In wild-type animals, hypoxia induced a rapid transient increase in FBP levels that gradually returned towards baseline. In contrast, *Ec*STH-expressing neurons exhibited a delayed and attenuated increase in FBP that reached a sustained plateau throughout hypoxia **(Figure 3B)**. These responses are consistent with elevated NADH/NAD⁺ imposing a constraint on glycolytic dynamics during energetic stress.

**Figure 3:**
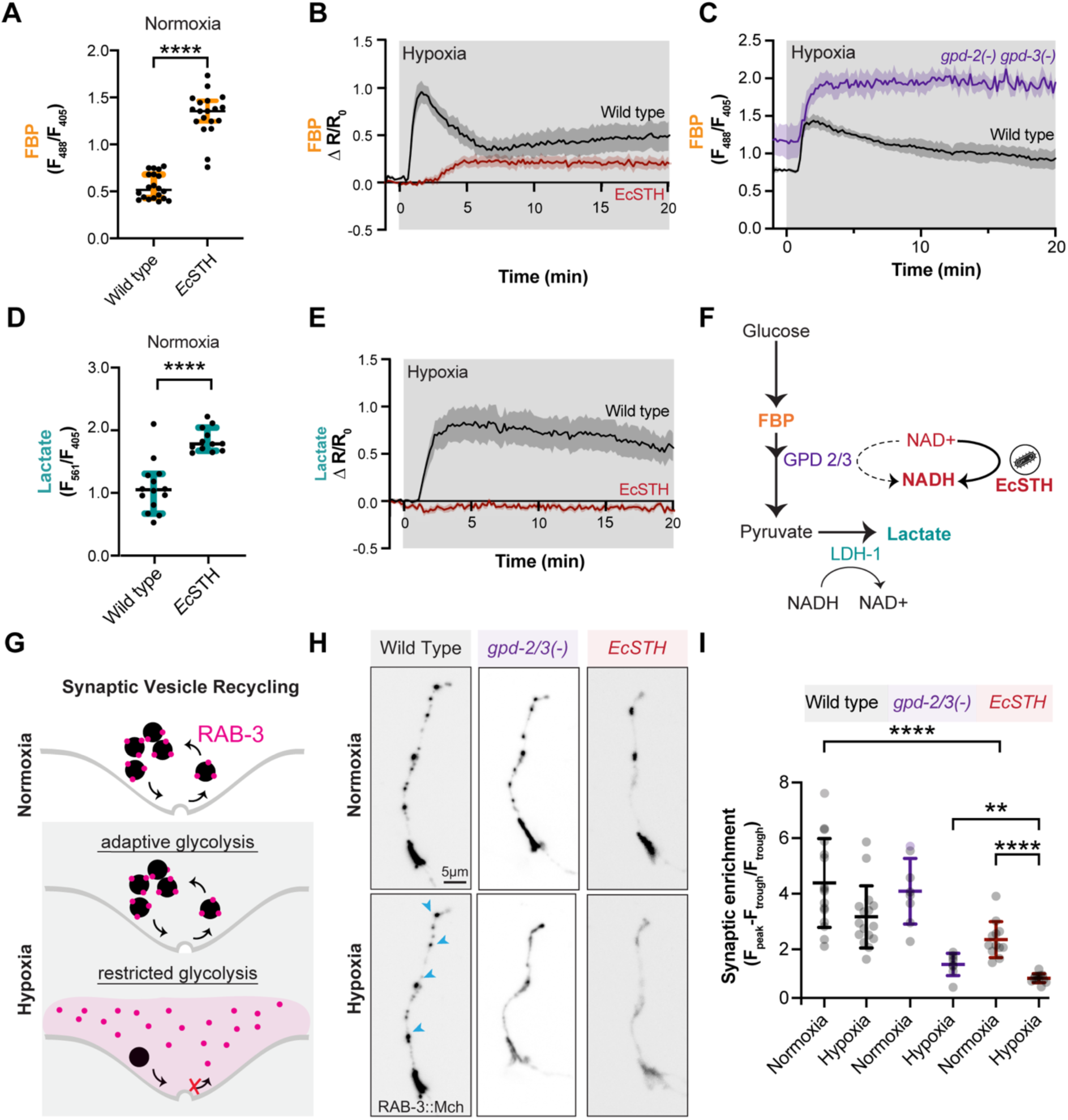
Elevated NADH/NAD⁺ ratios modify glycolytic dynamics and alter synaptic organization. **(A)** Quantification of steady-state FBP levels (F488/F405) in AIY neurons in wild-type and *Ec*STH-expressing animals. **(B)** Time course of relative FBP dynamics during 20 minutes of hypoxia, plotted as ΔR/R₀ (change in sensor ratio normalized to baseline), in wild-type (black) and *Ec*STH-expressing (maroon) animals. *Ec*STH expression attenuated the hypoxia-induced increase in FBP. **(C)** Time course of FBP dynamics (F488/F405) during hypoxia in wild-type (black) and *gpd-2(−) gpd-3(−)* animals (purple). Loss of *gpd-2* and *gpd-3* caused FBP accumulation during hypoxia. **(D)** Quantification of steady-state lactate levels measured using FiLa (F561/F405) in AIY neurons in wild-type and *Ec*STH-expressing animals. *Ec*STH expression increased baseline lactate levels relative to wild type. **(E)**Time course of relative lactate dynamics during 20 minutes of hypoxia, plotted as ΔR/R₀, in wild-type (black) and *Ec*STH-expressing (maroon) animals. *Ec*STH expression attenuated hypoxia-induced lactate accumulation. **(F)** Schematic illustrating proposed effects of altered NADH/NAD⁺ balance on glycolytic pathway behavior. Under normoxic conditions, increased NADH/NAD⁺ promotes lactate accumulation through LDH-1-mediated NADH recycling. During hypoxic stress, elevated NADH/NAD⁺ is proposed to constrain lower glycolysis at the NAD⁺-dependent GAPDH step encoded by *gpd-2* and *gpd-3*, resulting in accumulation of upstream glycolytic intermediates and reduced lactate production. **(G)** Schematic illustrating the relationship between glycolytic state and synaptic vesicle organization during hypoxia. Due to hypoxia-induced glycolytic upregulation, synaptic vesicle-associated RAB-3 remains enriched at presynaptic sites. In contrast, disruption of glycolysis impairs synaptic vesicle recycling, resulting in diffuse cytosolic localization of RAB-3. **(H)** Representative images of RAB-3::mCherry localization in AIY neurons under normoxia and hypoxia in wild-type, *gpd-2(−) gpd-3(−)*, and *Ec*STH-expressing animals. Arrowheads indicate punctate presynaptic localization. Scale bar, 5 μm. **(I)** Quantification of synaptic enrichment of RAB-3::mCherry in AIY neurons under normoxia and hypoxia. Synaptic enrichment was calculated as (Fpeak−Ftrough)/Ftrough: detailed in Methods. Hypoxia significantly reduced synaptic enrichment in *gpd-2(−) gpd-3(−)* and *Ec*STH-expressing animals relative to wild type. *Ec*STH-expressing animals also exhibited reduced synaptic enrichment under normoxic conditions compared to wild type. Each dot represents a single animal. Bars represent mean ± SD. Statistical comparisons were performed using an unpaired Student’s t test, except in (I), where paired Student’s t tests were used for within-genotype comparisons between normoxia and hypoxia. **** p < 0.0001, ** p < 0.01. Shaded regions in B, C and E represent mean ± SEM.

If FBP accumulation upon *Ec*STH expression reflects inhibition of lower glycolytic throughput at the GAPDH step, then direct genetic disruption of GAPDH should phenocopy this FBP accumulation. In *C. elegans*, GAPDH is encoded by four paralogs (*gpd-1, gpd-2, gpd-3 & gpd-4*). Published single-cell transcriptomics indicate that *gpd-2 and gpd-3* are the predominant isoforms expressed in AIY neurons^35^. We therefore generated a *gpd-2(−) gpd-3(−)* double mutant. Because *gpd-2* and *gpd-3* are co-transcribed as part of an operon, both genes were simultaneously deleted using CRISPR–Cas9 genome editing. Using HYlight to monitor FBP levels, we observed a hypoxia-induced increase and marked accumulation of FBP in *gpd-2(−) gpd-3(−)* mutants compared to wild-type animals **(Figure 3C)**. This accumulation and sustained elevation resembled the FBP dynamics observed upon *Ec*STH expression, supporting the interpretation that elevated FBP levels arise from impaired lower glycolysis and that redox imbalance is sufficient to constrain glycolytic throughput.

We next asked how elevated NADH/NAD⁺ influences lower glycolytic state by monitoring lactate dynamics using FiLa. Under baseline normoxic conditions, *Ec*STH expression significantly increased lactate levels relative to wild-type animals **(Figure 3D)**, indicating active NAD⁺ regeneration through LDH-1. However, during hypoxia, *Ec*STH-expressing neurons failed to exhibit the robust increase in lactate observed in wild-type animals **(Figure 3E)**. Thus, despite elevated baseline lactate levels, *Ec*STH expression blunted dynamic lactate responses during energetic stress. Together, the sustained accumulation of FBP and impaired lactate responses during hypoxia support a model in which elevated NADH/NAD⁺ constrains lower glycolysis, likely through limitation of the NAD⁺-dependent GAPDH reaction **(Figure 3F)**.

Previous studies have shown that glycolysis is required for synaptic vesicle recycling, particularly under conditions of mitochondrial inhibition or hypoxia^36–39^. We hypothesized that elevated NADH/NAD⁺ might impair the ability of neurons to maintain synaptic vesicle organization during energetic stress by altering the cellular glycolytic state. We therefore asked whether elevated NADH/NAD⁺ was sufficient to impair synaptic vesicle organization in a manner analogous to glycolytic mutants. Synaptic vesicles were visualized by tagging the presynaptic vesicle-associated protein RAB-3 with mCherry **(Figure 3G)**. In wild-type animals, RAB-3 localized to discrete puncta along synaptic zone 2 of AIY, corresponding to individual synaptic boutons^40^. Quantification of synaptic enrichment confirmed a marked reduction in punctate localization under hypoxia in *gpd-2(−) gpd-3(−)* mutants relative to wild type **(Figure 3H,I).** *Ec*STH expression produced an even stronger defect under hypoxia, with *Ec*STH-expressing neurons exhibiting markedly reduced synaptic enrichment, lower than that observed in *gpd-2(−) gpd-3(−)* mutants.

Unlike *gpd-2(−) gpd-3(−)* mutants, which retained near wild-type synaptic organization under normoxic conditions, *Ec*STH expression also modestly reduced synaptic enrichment in the presence of oxygen, despite synaptic puncta remaining qualitatively organized **(Figure 3H,I)**. The presence of a synaptic defect in normoxia in *Ec*STH-expressing neurons suggests that elevated NADH/NAD⁺ alters neuronal physiology through mechanisms beyond acute restriction of glycolysis alone. Under normoxic conditions, *Ec*STH expression increased both FBP and lactate levels **(Figures 3A,D)**, indicating that neurons remain metabolically active and continue to produce lactate to regenerate NAD⁺. This compensatory lactate production may help sustain glycolytic throughput despite elevated NADH/NAD⁺ ratios. However, increased conversion of pyruvate to lactate could simultaneously reduce pyruvate availability for mitochondrial metabolism, while elevated NADH/NAD⁺ may additionally perturb other NAD⁺-dependent metabolic processes required for maintenance of synaptic vesicle organization. Thus, although the broader metabolic state under normoxia remains difficult to resolve, these findings suggest that altered redox balance alone is sufficient to compromise synaptic organization even before the severe lower glycolytic restriction observed during hypoxia.

Together, these results demonstrate that elevated neuronal NADH/NAD⁺ produces a distinct glycolytic state characterized by elevated lactate production under normoxia, followed by sustained FBP accumulation, impaired lower glycolytic responses, and disrupted synaptic vesicle organization during transient hypoxia.

### Lowering NADH/NAD+ ratios increases glycolytic metabolites

We next asked whether facilitating NADH recycling to NAD⁺ through expression of *Lb*NOX could relieve redox constraints on glycolysis. We measured FBP using HYlight in the presence of *Lb*NOX and observed moderately increased baseline FBP levels **(Figure 4A)**. Endogenously, conversion of pyruvate to lactate regenerates NAD⁺; however, in the presence of LbNOX, pyruvate-to-lactate conversion is reduced **(Figure 4B)**, likely due to substrate competition for NADH^29^.

**Figure 4:**
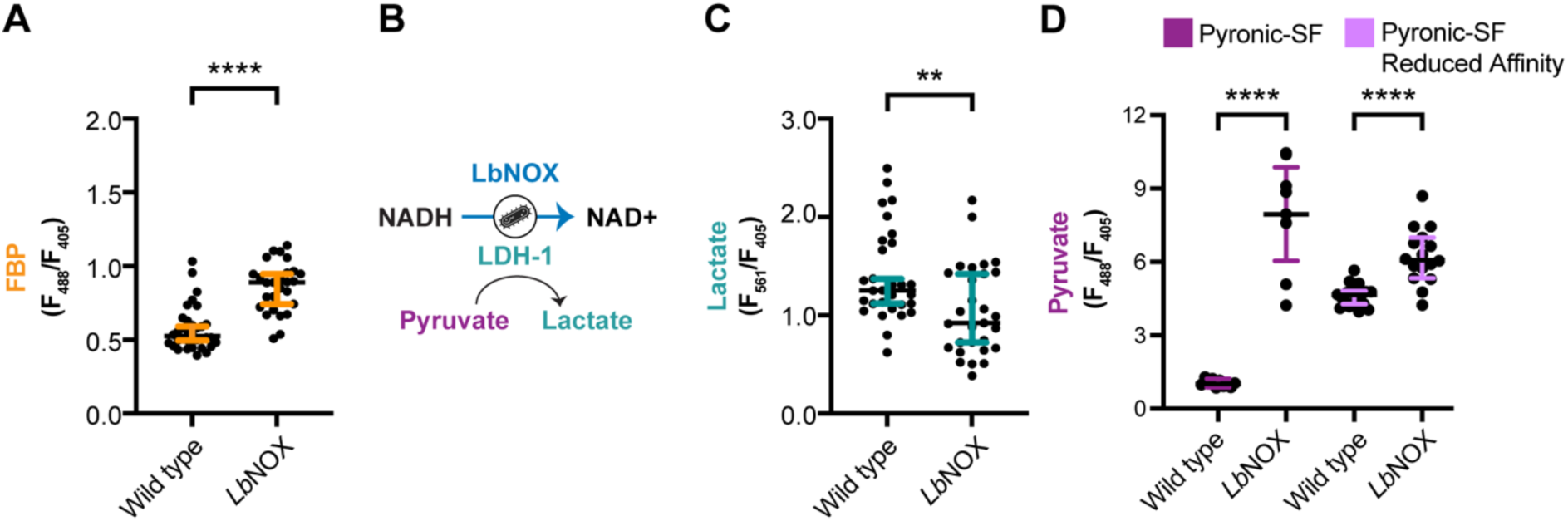
Lowering NADH/NAD+ ratios via LbNOX increases glycolytic metabolites. **(A)** Quantification of steady-state FBP levels (F488/F405) in AIY neurons in wild-type (black) and LbNOX-expressing (orange) animals. LbNOX expression increases FBP levels relative to wild type. Each dot represents a single animal. Bars represent mean ± 95% CI. **** p < 0.0001. **(B)** Schematic illustrating LbNOX-mediated oxidation of NADH to NAD⁺, providing an alternative NADH recycling pathway to LDH-1, which normally regenerates NAD⁺ during pyruvate-to-lactate conversion. **(C)** Quantification of steady-state lactate levels (F561/F405), teal) and **(D)** pyruvate levels measured using Pyronic-SF (F488/F405, right; magenta) in wild-type and LbNOX-expressing animals. The reduced-affinity Pyronic-SF control is shown in light purple. Each dot represents a single animal. Bars represent mean ± 95% CI. Statistical comparisons were performed using an unpaired Student’s t test. **** p < 0.0001; ** p < 0.01.

To assess downstream metabolites, we adapted the pyruvate sensor Pyronic-SF^16^ and expressed it in AIY neurons. *Lb*NOX expression resulted in significantly decreased lactate levels **(Figure 4C)** and increased pyruvate levels **(Figure 4D)**. A reduced-affinity version of Pyronic-SF showed markedly attenuated responses upon LbNOX expression, consistent with reduced pyruvate binding and supporting that the observed signal changes are specific to pyruvate (**Figure 4D**). Using this multisensor framework, *Lb*NOX shifts steady-state metabolites (increased FBP, increased pyruvate and reduced lactate), consistent with increased lower glycolytic throughput. Decreasing the NADH/NAD⁺ ratio could also alter other metabolic pathways, including mitochondrial energy production via NADH shuttling. To determine whether the observed changes reflected mitochondrial energetic stress, we examined synaptic vesicle localization using RAB-3. *Lb*NOX expression did not alter RAB-3 localization under normoxia, indicating no detectable defects in synaptic vesicle organization **(Figure S3A)**. HYlight responses during hypoxia were similar to wild type **(Figure S3B)**. Lactate levels were reduced at baseline but increased during hypoxia and remained elevated, potentially reflecting conversion of excess pyruvate to lactate under energetic stress. Interpretation of *Lb*NOX effects during hypoxia is complicated by its dependence on oxygen and proton availability, which may limit its activity under these conditions. Together, these results indicate that elevation of NADH/NAD⁺ inhibits lower glycolysis, whereas NADH oxidation by *Lb*NOX shifts metabolite pools consistent with relieved redox constraint.

### Genetic dissection of NADH buffering pathways reveals redox control of neuronal glycolytic dynamics

Having established that elevated NADH/NAD⁺ ratios inhibit glycolysis and synaptic vesicle recycling **(Figure 3)**, we next asked how neurons physiologically buffer NADH accumulation during hypoxic stress. Cytosolic NADH can be oxidized through both direct enzymatic pathways and mitochondrial shuttle systems. LDH-1 regenerates NAD⁺ through conversion of pyruvate to lactate, whereas GPDH-2 and MDH-1 function as cytosolic components of the glycerol-3-phosphate and malate–aspartate shuttles, respectively, which couple cytosolic redox state to mitochondrial metabolism **(Figure 5A)**. In some contexts, however, the glycerol-3-phosphate pathway can function independently of its mitochondrial counterpart, enabling cytosolic NAD⁺ regeneration^41,42^. To define the relative contribution of these pathways *in vivo*, we generated CRISPR-mediated null alleles of *ldh-1*, *gpdh-2*, and *mdh-1* **(Figure 5B)** and monitored NADH/NAD⁺, FBP, and lactate dynamics in AIY neurons during hypoxia.

**Figure 5.**
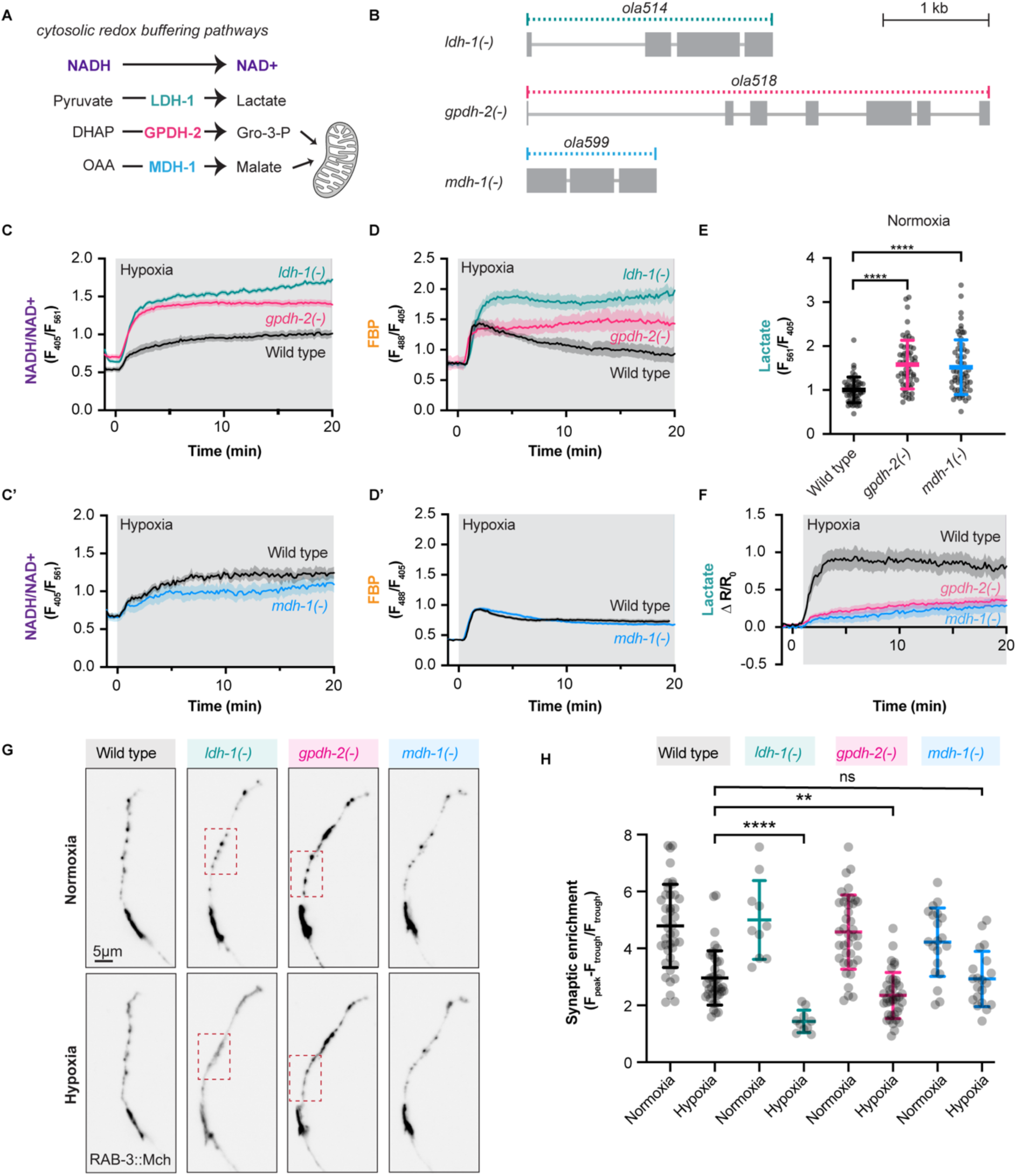
Distinct cytosolic NADH recycling pathways differentially maintain redox balance and shape glycolytic state in neurons. **(A)** Schematic of cytosolic NADH recycling pathways in neurons. LDH-1 oxidizes NADH during conversion of pyruvate to lactate. GPDH-2 oxidizes NADH during reduction of dihydroxyacetone phosphate (DHAP) to glycerol-3-phosphate. MDH-1 oxidizes NADH during reduction of oxaloacetate (OAA) to malate. **(B)** Schematic of gene structure and deletion alleles for ldh-1(ola514), *gpdh-2(ola518),* and *mdh-1(ola599).* Scale bar, 1 kb. **(C–D′)** Time course of NADH/NAD⁺ ratios measured with SoNar (C, C′) and FBP levels measured with HYlight (D, D′) during hypoxia in wild-type animals and the indicated mutants. Wild type, black; *ldh-1(-)*, teal; *gpdh-2(-)*, pink; *mdh-1(-)*, blue. Shaded regions represent mean ± SEM. **(E)** Baseline lactate levels measured with FiLa in wild-type animals and *gpdh-2(-)* and *mdh-1(-)* mutants under normoxic conditions. **(F)** Time course of lactate dynamics during hypoxia measured with FiLa and normalized to the baseline of each strain. Wild-type animals (black) exhibit robust lactate accumulation during hypoxia, whereas *gpdh-2(-)* (pink) and *mdh-1(-)* (blue) mutants show attenuated lactate responses. Shaded regions represent mean ± SEM **(G)** Representative images of RAB-3::mCherry localization in AIY neurons under normoxia and hypoxia in wild-type, *ldh-1(-)*, *gpdh-2(-)*, and *mdh-1(-)* animals. Scale bar, 5 µm. **(H)** Quantification of synaptic enrichment of RAB-3::mCherry in AIY neurons under normoxia and hypoxia, calculated as (Fpeak−Ftrough)/Ftrough. *ldh-1(-)* and *gpdh-2(-)* animals exhibit reduced synaptic enrichment during hypoxia, whereas *mdh-1(-)* animals are similar to wild type. Statistical comparisons were performed using an unpaired Student’s t test, except in (H), where paired Student’s t tests were used for within-genotype comparisons between normoxia and hypoxia. Each dot represents one animal. Bars represent mean ± SD. ns, not significant; ****P < 0.0001.

Loss of LDH-1 produced the strongest defect in redox homeostasis. During hypoxia, *ldh-1(-)* neurons exhibited a pronounced elevation in NADH/NAD⁺ ratios **(Figure 5C)** together with marked accumulation of FBP **(Figure 5D)**, consistent with impaired NAD⁺ regeneration and reduced glycolytic throughput, supporting the conclusion that LDH-1 is the dominant route for cytosolic NADH recycling during acute hypoxic stress.

Loss of GPDH-2 also impaired redox balance and glycolytic state, but less severely than loss of LDH-1. *gpdh-2(-)* neurons showed a clear increase in NADH/NAD⁺ during hypoxia, accompanied by accumulation of FBP, although both phenotypes were attenuated relative to *ldh-1(-)* animals **(Figure 5C,D)**. These results indicate that GPDH-2 contributes to cytosolic NADH recycling in neurons during hypoxia but plays a secondary role relative to LDH-1. In parallel, *gpdh-2(-)* mutants exhibited elevated baseline lactate levels **(Figure 5E)** and a blunted hypoxia-induced lactate response **(Figure 5F)**, consistent with increased reliance on LDH-1-mediated NADH recycling under basal conditions and reduced dynamic capacity during stress. In contrast, loss of MDH-1 had little effect on NADH/NAD⁺ **(Figure 5C’)** or FBP dynamics **(Figure 5D’)** during hypoxia. *mdh-1(-)* neurons were largely indistinguishable from wild type for both readouts, indicating that MDH-1 makes minimal contribution to maintaining cytosolic redox balance or sustaining glycolytic flux under these conditions. As in *gpdh-2(-)* mutants, however, lactate levels were elevated at baseline **(Figure 5E)** and showed reduced further increase during hypoxia **(Figure 5F)**, suggesting compensatory engagement of lactate production without major disruption of glycolytic state.

Across all mutants, the magnitude of FBP accumulation tracked closely with the magnitude of NADH/NAD⁺ elevation, arguing that graded impairment of cytosolic NAD⁺ regeneration produces graded inhibition of glycolytic flux. Together, these data support a hierarchical organization of NADH recycling pathways in AIY neurons, with LDH-1 serving as the dominant physiological buffer, GPDH-2 providing a substantial but smaller contribution, and MDH-1 playing little detectable role under these conditions.

To determine whether graded disruption of NADH recycling pathways also produces functional consequences, we next examined synaptic vesicle organization using the presynaptic marker RAB-3::mCherry **(Figure 5G,H)**. Consistent with the severe metabolic defects observed in *ldh-1(-)* mutant animals, hypoxia produced a pronounced reduction in synaptic enrichment in *ldh-1(-)* mutants relative to wild type. *gpdh-2(-)* mutants also exhibited impaired synaptic enrichment during hypoxia, although the defect was less severe than that observed in *ldh-1(-)* animals, supporting a graded relationship between disruption of cytosolic NADH recycling, glycolytic impairment, and synaptic vesicle recycling. In contrast, *mdh-1(-)* animals remained largely similar to wild type under both normoxic and hypoxic conditions.

To further test the relationship between these pathways, we examined genetic interactions between LDH-1 and GPDH-2 **(Figure 6A).** Loss of *gpdh-2* in an *ldh-1(-)* background did not substantially increase NADH/NAD⁺ accumulation or FBP levels beyond those observed in *ldh-1(-)* animals alone during hypoxia **(Figure 6B,C)**. Notably, *ldh-1(-)* mutants exhibited elevated FBP levels comparable to those observed upon genetic disruption of the glycolytic enzyme GAPDH (*gpd-2(-) gpd-3(-)*), indicating that loss of LDH-1 is sufficient to impose a near-maximal constraint on glycolytic flux. Thus, once LDH-1-dependent NADH recycling is lost, removing GPDH-2 does not produce a strong additional defect, indicating that GPDH-2 is not able to compensate for LDH-1 loss at endogenous expression levels.

**Figure 6:**
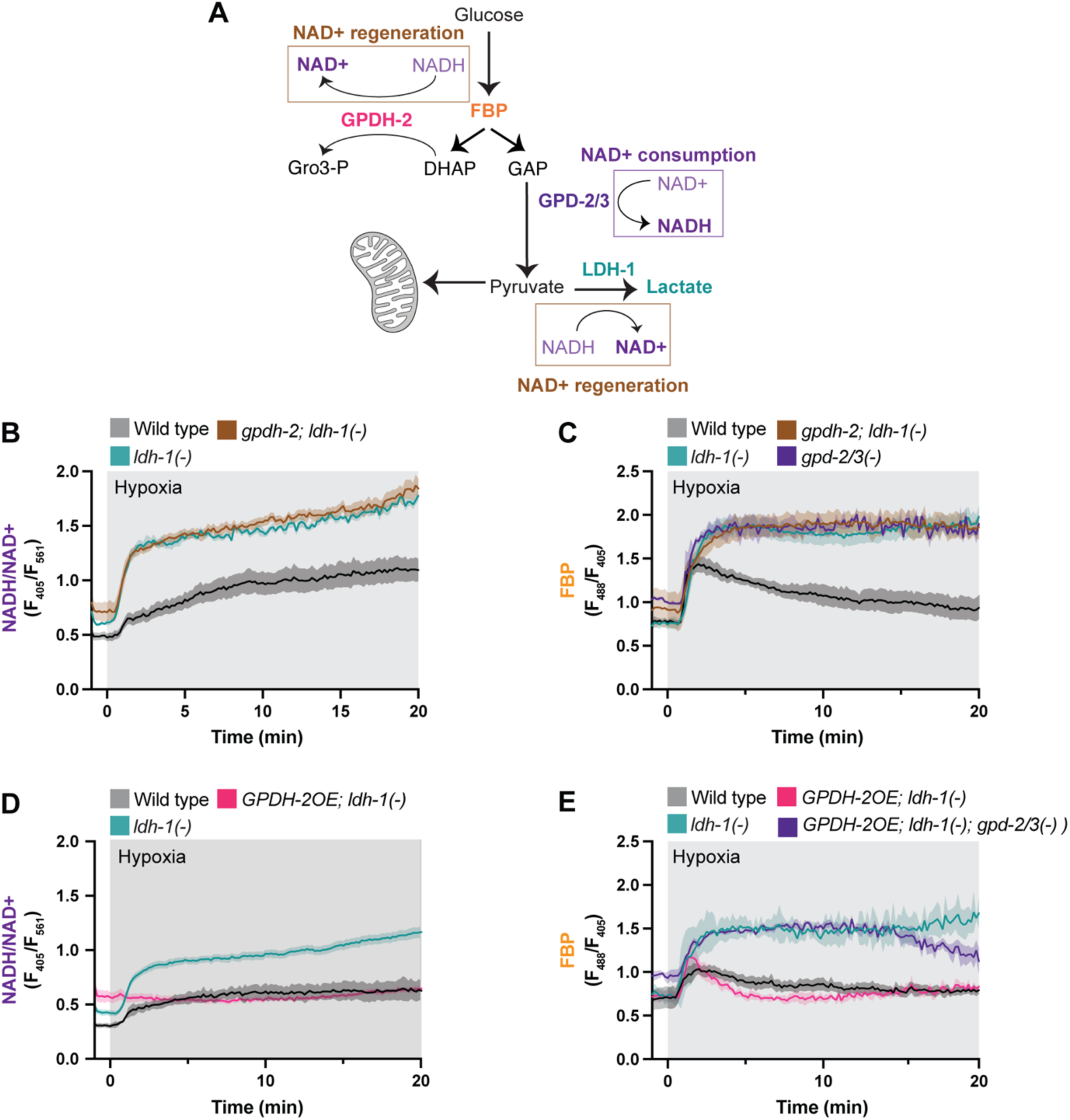
GPDH-2 overexpression compensates for loss of LDH-1 by restoring NADH/NAD⁺ balance and FBP consumption. **(A)** Schematic illustrating the relationship between glycolytic intermediates and cytosolic NADH/NAD⁺ balance. NAD⁺ is consumed during the GAPDH reaction (GPD-2/3) and regenerated through LDH-1-mediated conversion of pyruvate to lactate or GPDH-2-mediated conversion of dihydroxyacetone phosphate (DHAP) to glycerol-3-phosphate. The schematic highlights the pathways examined to assess compensatory mechanisms of cytosolic NAD⁺ regeneration. **(B)** NADH/NAD⁺ ratios (F405/F561) during hypoxia in wild-type (black), *ldh-1(−)* (teal), and *gpdh-2(−); ldh-1(−)* (brown) animals. **(C)** FBP levels (F488/F405) during hypoxia in wild-type (black), *ldh-1(−)* (teal), *gpdh-2(−); ldh-1(−)* (brown), and *gpd-2(−) gpd-3(−)* (purple) animals. **(D)** NADH/NAD⁺ ratios during hypoxia in wild-type animals (black), *ldh-1(−)* mutants (teal), and GPDH-2 overexpression in the *ldh-1(−)* background (pink). **(E)** FBP levels during hypoxia in wild-type animals (black), ldh-1(−) mutants (teal), GPDH-2 overexpression in an *ldh-1(−)* background (pink), and GPDH-2 overexpression in a *ldh-1(−); gpd-2(−); gpd-3(−)* triple mutant background (purple). Shaded regions in (B–E) represent mean ± SEM.

We next asked whether GPDH-2 retains the capacity to support NADH recycling when expressed at higher levels. Overexpression of GPDH-2 in *ldh-1(-)* neurons markedly attenuated the hypoxia-induced increase in NADH/NAD⁺ **(Figure 6D)** and restored FBP dynamics toward wild-type levels **(Figure 6E)**, demonstrating that GPDH-2 can substitute for LDH-1 when sufficiently expressed. This suppression was abolished in animals lacking *gpd-2* and *gpd-3*, indicating that GPDH-2–mediated NAD⁺ regeneration restores flux through the NAD⁺-dependent GAPDH step and enables FBP consumption via lower glycolysis rather than by diversion of intermediates into glycerol-3-phosphate.

Together, these findings reveal a hierarchical organization of neuronal NADH recycling pathways, with LDH-1 providing the dominant capacity for NAD⁺ regeneration and GPDH-2 contributing a smaller, secondary source of NAD⁺ regeneration. While GPDH-2 is required to support redox balance and glycolysis in neurons, it is not sufficient to buffer the acute NADH accumulation induced by loss of LDH-1. However, when upregulated, GPDH-2 retains the capacity to restore redox balance and thereby sustain glycolysis.

## DISCUSSION

### Integrated biosensor approaches enable quantitative interpretation of glycolysis in vivo

Metabolism is both dynamic and spatially compartmentalized, requiring approaches that move beyond isolated metabolite measurements toward integrated interpretation of pathway state *in vivo*. Here, we establish a multisensor framework to interpret glycolytic dynamics in intact neurons by monitoring an upstream intermediate, downstream metabolites, and cytosolic redox state. No single metabolite provides a sufficient proxy for glycolytic flux. Lactate, for example, can reflect glycolytic output but may also derive from extracellular uptake or intercellular exchange, complicating its interpretation as a cell-autonomous readout. Similarly, accumulation of the upper glycolytic intermediate FBP can reflect either increased glycolytic input or impaired flux through lower glycolysis, as observed under conditions of elevated NADH/NAD⁺. Redox state itself is also ambiguous when considered in isolation, as shifts in NADH/NAD⁺ can arise from multiple metabolic processes. By comparing measurements of FBP, lactate, pyruvate, and redox state in the same identified neuron exposed to the same metabolic perturbation, our approach enables semi-quantitative inference of pathway behavior *in vivo*.

Building on our multisensor approach, accurate interpretation of biosensor measurements *in vivo* requires careful consideration of both sensor properties and biological context. First, binding-deficient controls are essential to distinguish metabolite-dependent signals from non-specific fluorescence changes. Second, where possible, genetic mutants provide important *in vivo* validation by linking sensor responses to defined metabolic steps, while orthogonal genetic tools such as heterologous enzymes that modulate metabolic state^27^ can provide complementary validation when genetic approaches are limited. Third, sensor readouts can be influenced by environmental factors such as pH, necessitating independent validation of these effects. Fourth, ratiometric sensor designs are particularly important *in vivo*, because they help correct for variability in expression level and animal movement, but require careful assessment of photobleaching, especially in sensors that rely on multiple fluorophores. Finally, because single-metabolite measurements are inherently ambiguous, our results highlight that sensor signals must be interpreted in the context of their pathway to infer pathway-level metabolic dynamics. Future advances in biosensor engineering and spectral multiplexing may ultimately enable simultaneous measurements of multiple metabolites within single cells.

### Cytosolic redox state directly constrains glycolysis and neuronal function

Beyond serving as a readout, cytosolic redox state has been proposed to directly regulate glycolytic flux^20,22,23,26^. Studies in proliferating cells have shown that NAD⁺ demand, rather than ATP demand, can be a primary driver of aerobic glycolysis^20,23^, and that lactate accumulation can limit glycolytic flux through redox-dependent mechanisms in T cells^22^. However, whether redox state causally constrains glycolysis *in vivo* has remained unclear, as prior studies have largely examined redox effects in the context of chronic perturbations such as hypoxia^43^, alcohol exposure^44^, mitochondrial inhibition^45,46^, or metabolic disease^30,47^, where shifts in NADH/NAD⁺ are typically inferred indirectly and alongside widespread metabolic changes that complicate causal interpretations. By directly manipulating neuronal redox state using heterologous enzymes, we demonstrate that NADH/NAD⁺ elevation is sufficient to constrain glycolysis *in vivo*. Importantly, these metabolic changes are accompanied by alterations in synaptic vesicle organization, extending the redox-glycolysis relationship established in cultured cells to intact neurons and establishing cytosolic redox balance as an active determinant of both glycolytic flux and synaptic function.

### NAD⁺ regeneration pathways are not interchangeable and differ in their capacity to sustain glycolysis

We find that cytosolic NADH recycling pathways are not functionally redundant but instead differ in their capacity to regenerate NAD⁺. Both LDH-1 and GPDH-2 contribute to cytosolic NADH recycling, consistent with prior work demonstrating cooperative and partially compensatory roles for these pathways^49^. However, rather than acting as interchangeable buffers, these pathways operate within a capacity-limited framework. LDH-1 provides the dominant route for NAD⁺ regeneration during acute stress, whereas GPDH-2 plays an auxiliary role. Consequently, loss of LDH-1 imposes a near-maximal constraint on lower glycolysis that endogenous GPDH-2 cannot fully compensate for. Nevertheless, the increase in NADH/NAD⁺ observed in *gpdh-2(-)* mutants indicates that GPDH-2 contributes meaningfully to cytosolic redox balance under physiological conditions. The ability of GPDH-2 overexpression to restore redox balance and glycolytic flux further demonstrates that this pathway retains latent capacity that is not fully utilized under normal conditions.

These differences reflect the metabolic context in which NAD⁺ regeneration occurs. LDH-1 regenerates NAD⁺ while preserving carbon within glycolysis through conversion of pyruvate to lactate, thereby supporting continued flux through lower glycolysis. In contrast, GPDH-2 couples NADH oxidation to the reduction of dihydroxyacetone phosphate, diverting carbon into glycerol-3-phosphate and downstream lipid metabolism. Thus, engagement of GPDH-2 introduces a trade-off between maintaining redox balance and preserving glycolytic throughput versus redirecting carbon into alternative metabolic fates^42,48^.

Reductive stress is increasingly associated with conditions such as ischemia^50–52^, mitochondrial dysfunction^41,45,53^, overnutrition^54,55^, metabolic disease^30,47^ and aging^56,57^. Our findings suggest that elevated NADH/NAD⁺ can reshape glycolytic dynamics, with consequences for synaptic function and vulnerability to energetic stress. Extending this framework across neuronal types and disease models may help distinguish when shifts in redox state are consequences of altered metabolism and when they become causal constraints on glycolytic function and neuronal physiology.

## ACKNOWLEDGMENTS

We thank the Yang lab (East China University of Science and Technology, Shanghai, China) for sharing the sequences of SoNaR and FiLa. We thank the Hammarlund lab (Yale University, New Haven CT, USA) for sharing the codon-optimized construct of R-iLACCO1.2. We thank the Keck Oligo Synthesis Resource at Yale for their assistance with synthesis of short single-stranded DNA oligos. We also thank Gary Yellen, Richard Goodman and members of the Colón-Ramos lab for their thoughtful comments and discussions related to this project. This work was supported by National Institutes of Health grants to D.C.-R. (R35NS132156 and R01NS076558).

## METHODS

### Strains and oligonucleotides used

All strains (listed in Supplementary Table S1) were cultured at 20◦C on Nematode Growth Medium (NGM) plates seeded with 100ul of OP50 and mutant combinations were generated using standard methods^58^. Reference alleles indicated as *gene(-)* are as follows: *pfk-1.1(ola458), ldh-1(ola514)*, *gpd-2 gpd-3(ola542)*, *gpdh-2(ola518)* and *mdh-1(ola599)*. Sequences of oligonucleotides used to genotype different mutant combinations are in Supplementary Table S3 (*pfk-1.1*: P01-P03, *ldh-1*: P04-P06, *gpd-2 gpd-3*: P07-P09, *gpdh-2*: P10-P12 and *mdh-1*: P13-P15). \

### Transgenesis and Genome Editing

Transgenic *Caenorhabditis elegans* strains were generated by standard germline injection techniques^59^. Plasmids were assembled using Gibson Assembly. Primers were designed in SnapGene (version 7.0.3) to amplify the desired vector backbone and insert sequences. DNA fragments were PCR-amplified using CloneAmp HiFi PCR Premix, purified, and assembled using the Gibson Assembly 2× Enzyme Premix according to the manufacturer’s instructions. Assembly reactions were transformed into Stellar Competent Cells, and transformants were selected on LB agar plates containing ampicillin. All plasmids were purified from bacterial culture using QIAprep Spin Miniprep Kit (QIAGEN). All plasmid constructs were verified by Sanger sequencing or whole plasmid sequencing was performed by Plasmidosaurus using Oxford Nanopore Technology with custom analysis and annotation. A complete list of plasmids is provided in Supplementary Table 2.

Transgenic animals were generated by expressing plasmids as extrachromosomal or integrated arrays. The plasmids DACR2436, which drives mCherry expression in the intestine, and DACR1708, which drives GFP expression in coelomocytes, were used as co-injection markers to identify transgenic animals.

Gene deletions of *ldh-1*, *gpdh-2*, *mdh-1*, *pfk-1.1* and *gpd-2 & gpd-3* were made by CRISPR by injecting Cas9 (PNA Bio), crRNA, tracrRNA (Horizon Discovery) and an oligonucleotide homology repair template^60^ (Keck Oligo synthesis resource at Yale), into the *C. elegans* distal gonad. Screening for plates with successfully edited F1 animals was performed using the *dpy-10* co-conversion strategy^61^. Worms from the F1 generation with the *roller* phenotype were singled out and sequenced for the deletion and the resulting homozygous progeny were outcrossed two times prior to use. For each gene, two crRNAs and one single stranded homology repair template were used (Supplementary Table S2: *pfk-1.1*: P16-P18, *ldh-1*: P19-P21, *gpd-2 gpd-3*: P22-P24, *gpdh-2*: P25-P27 and *mdh-1*: P28-P30).

### Design and validation of metabolic biosensors and redox-modulating enzymes

The coding sequences for the biosensors SoNar, FiLa, R-iLACCO1.2, and Pyronic-SF, as well as the *Escherichia coli* soluble transhydrogenase EcSTH and the *Lactobacillus brevis* water-forming NADH oxidase LbNOX, were synthesized as GeneArt Strings (Thermo Fisher Scientific). All sequences were codon optimized for *C. elegans* and engineered to contain a single synthetic intron within the first 100 bp to improve expression^62^.

SoNar and FiLa were cloned in frame with a T2A self-cleaving peptide followed by mScarlet-I3, enabling ratiometric measurements by normalizing sensor fluorescence to mScarlet-I3 fluorescence. R-iLACCO1.2 was cloned upstream of a T2A peptide and TagBFP2 to enable ratiometric measurements relative to BFP2 fluorescence. The design and optimization of the HYlight biosensor have been described previously^5^, and the same construct and base strain (DCR9089) were used here for subsequent genetic crosses.

All biosensor and enzyme constructs were expressed as extrachromosomal arrays and identified by co-injection of a fluorescent marker. To combine redox perturbations with metabolic reporters, EcSTH- and LbNOX-expressing strains were crossed into the corresponding biosensor backgrounds, generating animals carrying both the enzyme and sensor transgenes.

### Microfluidics and Mounting

For hypoxia imaging, worms were imaged using a PDMS microfluidic device that enabled precision gas flow of nitrogen (hypoxia) or air gas (normoxia), as previously described^5,36^. 6% agarose pads (in water) were affixed to the top of the PDMS devices where the gas exchange occurred. A 3 μL drop of 10 mM levamisole in M9, was placed onto the pad. Worms were transferred to the center of the drop, and a 22x22mm No. 1.5 glass coverslip was gently placed onto the pad.

### Confocal Microscopy

Live imaging was performed on a Nikon Ti2 microscope equipped with a CSU-W1 spinning disk confocal unit (Yokogawa) and a Hamamatsu Orca-Fusion BT CMOS camera. After mounting a slide or microfluidic device, a 5-min equilibration period was observed to allow the slide to thermally equilibrate with the slightly elevated ambient temperature on the microscope (with oxygen air flow). Images were acquired at 16-bit pixel depth using 405-, 488-, and 561-nm laser excitation as required.

All biosensor time-lapse imaging experiments were performed using a 10× objective unless otherwise indicated. Imaging settings were calibrated for different objectives to give an average baseline ratio of ∼0.50. Time-lapse recordings were acquired at 0.2–1 Hz for 20 min. Exposure times ranged from 100–200 ms depending on the biosensor.

For SoNar imaging, excitation was performed using 405 nm (28% laser power) and 561 nm (2% laser power) illumination. NADH/NAD⁺ levels were quantified using the F405/F561 ratio. For FiLa imaging, excitation was performed using 405 nm (20% laser power) and 561 nm (1% laser power) illumination. Lactate levels were quantified as (F561/F405). For HYlight imaging, excitation was alternated between 488 nm and 405 nm without changing the emission filter. Laser power was set to 8% and 2%, respectively. Fructose-1,6-bisphosphate levels were quantified using the F488/F405 ratio. For Pyronic-SF imaging, excitation was performed using 488 nm (5% laser power) and 405-nm (4% laser power) illumination and quantified using the F488/F405 ratio. For R-iLACCO1.2 imaging, excitation was performed using 405 nm and 561 nm illumination and quantified using the F561/F405 ratio.

High-magnification representative ratiometric images were acquired using a 60× objective. For analysis of synaptic vesicle organization, z-stack images were acquired using a 60× objective with 0.3-µm step spacing spanning the entire AIY neurite nearest the coverslip. Maximum-intensity projections were generated for analysis and display.

### Image processing and analysis

All image processing and analysis were performed using Fiji/ImageJ and GraphPad Prism 10. For biosensor imaging, images were background-subtracted using a 50-pixel rolling-ball algorithm. Regions of interest (ROIs) encompassing neuronal cell bodies were manually defined at selected frames throughout each recording. An ImageJ macro was used to linearly interpolate ROI positions between manually annotated frames, generating ROIs for all intermediate frames and compensating for minor movement during imaging. Mean fluorescence intensities were extracted from each channel for every frame.

Sensor ratios were calculated as F488/F405 (HYlight and Pyronic-SF), F405/F561 (SoNar), 1/(F405/F561) (FiLa), and F561/F405 (R-iLACCO1.2). For FiLa, the inverse ratio was reported such that increases in signal corresponded to increases in intracellular lactate. Photobleaching correction was performed using decay curves obtained from unstimulated control recordings and applied prior to ratio calculations.

For representative images, ratio images were generated using the Fiji Image Calculator and displayed using the mpl_magma lookup table. Ratio ranges were adjusted linearly as indicated in individual figures.

Synaptic vesicle organization was assessed by imaging mCherry::RAB-3 in AIY neurons as previously described^36^. Maximum-intensity projections were generated from z-stack images spanning the entire AIY neurite nearest the coverslip. Fluorescence intensity profiles were extracted from the Zone 3 region of the AIY neurite using the segmented line tool in Fiji/ImageJ. Synaptic enrichment was quantified using a custom sliding-window analysis adapted from previous ΔF/F-based measurements^36^. Briefly, local fluorescence maxima and minima were identified along the neurite within a 2-µm sliding window and used to calculate the relative enrichment of RAB-3 at presynaptic puncta:

### Photobleaching correction

To quantify photobleaching, animals were mounted on agarose pads on glass slides and sealed with nail polish to prevent drift associated with pad drying. For each sensor and excitation channel, fluorescence traces were extracted and normalized to the baseline value at t = −1 min (F/F₀ = 1). For each technical replicate, mean decay profiles were computed by averaging normalized traces across animals. To estimate bleaching rates, linear models were fit to the mean decay trace of each technical replicate for each excitation channel. Fits were constrained through the normalization point (t = −1 min, F/F₀ = 1) to maintain consistency with baseline normalization. The resulting slopes provided an estimate of channel-specific bleaching rates. Slopes from 3 independent technical replicates were combined by taking the median value for each channel. For iNapc/SoNar, corrections were applied independently to each channel to account for differential bleaching. For the reduced-affinity control FiLa-C, fluorescence excited at 405 nm exhibited negligible bleaching under these conditions, whereas the 561 nm channel showed measurable decay; accordingly, bleaching correction was applied only to the 561 nm channel for FiLa and FiLa-C.

### Statistics

Statistical analyses were performed using GraphPad Prism 10. Individual data points represent biologically independent animals. Experiments were independently repeated at least two times with similar results.

Comparisons between two groups were performed using two-tailed Student’s t-tests. Comparisons involving three or more groups were analyzed using one-way ANOVA followed by multiple-comparisons testing. Paired measurements obtained from the same animals under normoxic and hypoxic conditions were analyzed using paired t-tests. Exact statistical tests, sample sizes, and P values are reported in the corresponding figure legends.

Time-course data are presented as mean ± SEM. Scatter plots are presented as mean ± SD or mean ± 95% confidence interval as indicated in the figure legends. Statistical significance was defined as P < 0.05.

**Supplementary Table S1: STRAIN LIST**

**Supplementary Table S2:PLASMID LIST**

**Supplementary Table S3: OLIGONUCLEOTIDE LIST**

---

**Supplemental Figure S1.**
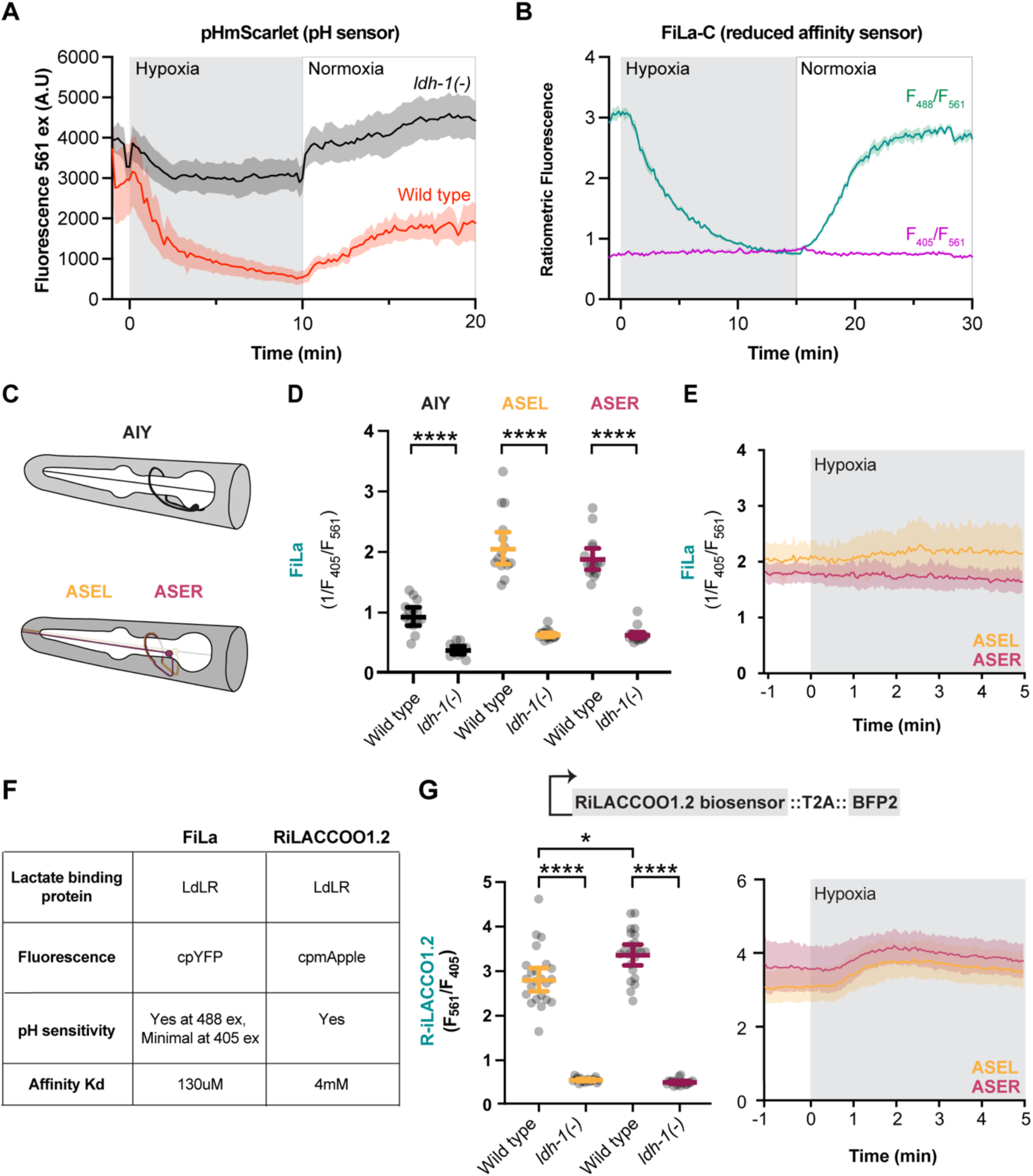
pH sensitivity and dynamic range of lactate biosensors FiLa and R-iLACCO1.2. **(A)** Cytosolic pH measured using pHmScarlet in wild-type (red) and *ldh-1(-)* (gray) animals during transitions between hypoxia and normoxia. **(B)** Ratiometric fluorescence of the reduced-affinity control sensor FiLa-C during hypoxia and reoxygenation. Fluorescence ratios were calculated as F488/F561 (green) and F405/F561 (purple). FiLa-C exhibited stimulus-dependent changes under 488 nm excitation but minimal changes under 405 nm excitation. **(C)** Schematic of AIY, ASEL, and ASER neurons used for lactate imaging experiments. **(D)** Quantification of baseline lactate levels measured using FiLa (F561/F405) in AIY, ASEL, and ASER neurons in wild-type and ldh-1(-) animals. **(E)** Time course of lactate dynamics measured using FiLa (F561/F405) during hypoxia in ASEL (orange) and ASER (pink) neurons. Shaded regions represent mean ± SEM. **(F)** Comparison of lactate biosensors FiLa and R-iLACCO1.2, including lactate-binding domain, fluorescent scaffold, pH sensitivity, and affinity (Kd). **(G)** Quantification of baseline lactate levels (*Left)* and hypoxia-induced lactate dynamics (*Right*) measured using the lower-affinity biosensor R-iLACCO1.2::T2A::BFP2 in ASEL (orange) and ASER (pink) neurons of wild-type and ldh-1(−) animals. Fluorescence ratios were calculated as F561/F405. Each dot in S1D and S1G represents a single animal. Bars represent mean ± 95% CI. Shaded regions represent mean ± SEM. **** p < 0.0001; * p < 0.05.

**Supplemental Figure S2.**
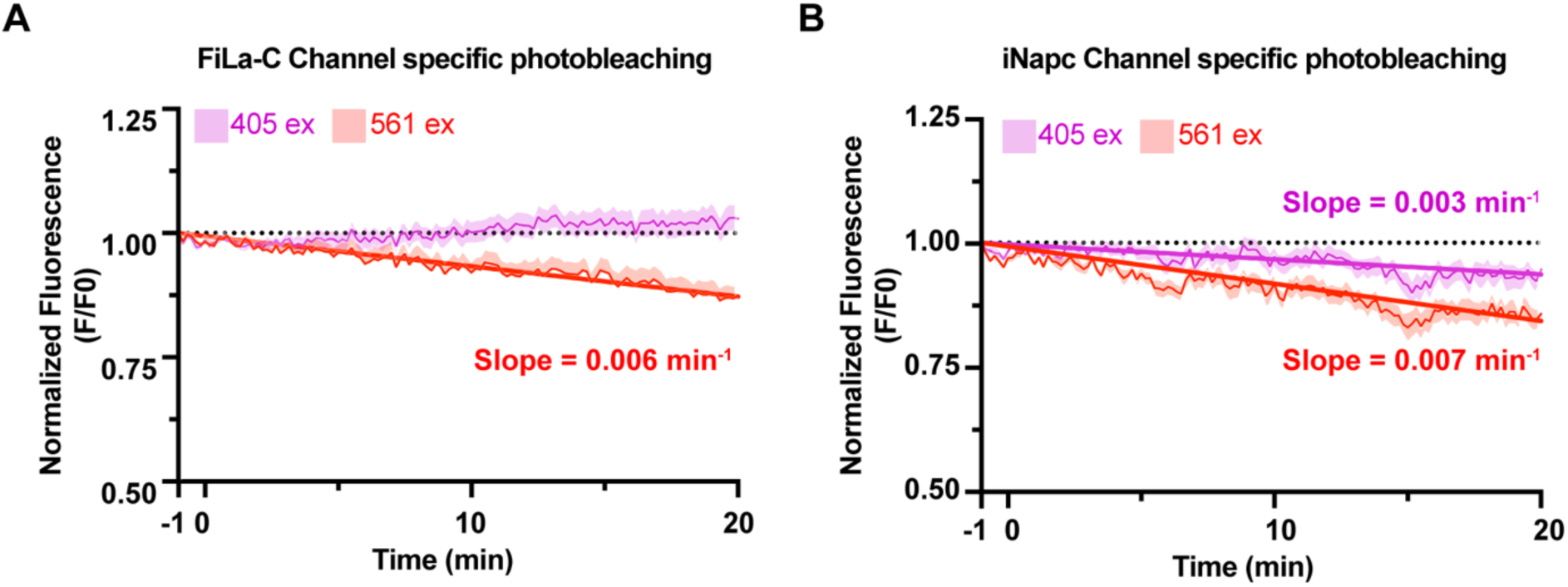
Characterization of channel-specific photobleaching in FiLa-C and iNAPc. **(A)** Channel-specific photobleaching of FiLa-C under 20 minute imaging conditions, shown as normalized fluorescence (F/F₀) for the 405 nm (purple) and 561 nm (red) channels. Linear fits were used to estimate bleaching rates (slope, min⁻¹), representing the fractional loss of fluorescence per minute. Bleaching of the 561 nm channel represents the primary source of ratio drift. **(B)** Channel-specific photobleaching of iNapc under 20 minute imaging conditions, shown as normalized fluorescence (F/F₀) for the 405 nm (purple) and 561 nm (red) channels. Linear fits indicate differential bleaching rates across channels, and photobleaching-induced drift in both channels was quantified and corrected prior to ratiometric analysis.

**Supplemental Figure S3.**
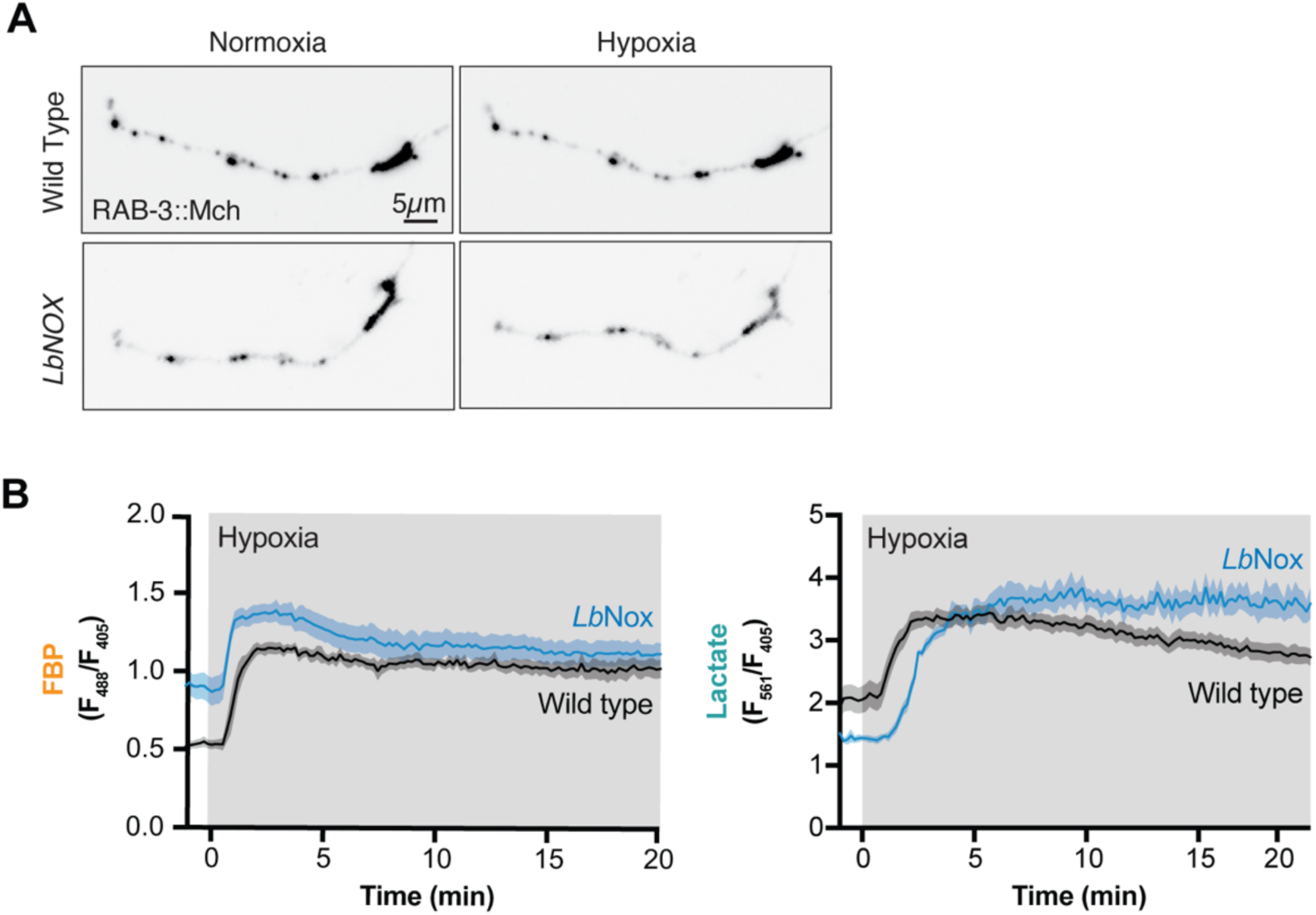
LbNOX expression does not disrupt RAB-3 synaptic localization under normoxic or hypoxic conditions. **(A)** Representative images of RAB-3::mCherry localization in AIY neurons under normoxia and hypoxia in wild-type and LbNOX-expressing animals. Scale bar, 5 μm. **(B)** Time course of FBP levels (F488/F405, left; orange) and lactate levels (1/(F405/F561), right; teal) during hypoxia in wild-type (black) and LbNOX-expressing (blue) animals. Shaded regions represent mean ± SEM.

## REFERENCES

1. Mink, J. W., Blumenschine, R. J. & Adams, D. B. Ratio of central nervous system to body metabolism in vertebrates: its constancy and functional basis. *American Journal of Physiology-Regulatory*, Integrative and Comparative Physiology 241, R203–R212 (1981).

2. Harris, J. J., Jolivet, R. & Attwell, D. Synaptic Energy Use and Supply. Neuron 75, 762–777 (2012).

3. Attwell, D. & Laughlin, S. B. An Energy Budget for Signaling in the Grey Matter of the Brain. J Cereb Blood Flow Metab 21, 1133–1145 (2001).

4. Díaz-García, C. M. et al. Neuronal Stimulation Triggers Neuronal Glycolysis and Not Lactate Uptake. Cell Metabolism 26, 361–374.e4 (2017).

5. Wolfe, A. D. et al. Local and dynamic regulation of neuronal glycolysis in vivo. Proc. Natl. Acad. Sci. U.S.A. 121, e2314699121 (2024).

6. Li, H. et al. Neurons require glucose uptake and glycolysis in vivo. Cell Reports 42, 112335 (2023).

7. Logothetis, N. K., Pauls, J., Augath, M., Trinath, T. & Oeltermann, A. Neurophysiological investigation of the basis of the fMRI signal. (2001).

8. Caprioli, R. M., Farmer, T. B. & Gile, J. Molecular Imaging of Biological Samples: Localization of Peptides and Proteins Using MALDI-TOF MS. Anal. Chem. 69, 4751–4760 (1997).

9. Wang, L. et al. Spatially resolved isotope tracing reveals tissue metabolic activity. Nat Methods 19, 223–230 (2022).

10. Miura, D. et al. Ultrahighly Sensitive in Situ Metabolomic Imaging for Visualizing Spatiotemporal Metabolic Behaviors. Anal. Chem. 82, 9789–9796 (2010).

11. Blacker, T. S. et al. Separating NADH and NADPH fluorescence in live cells and tissues using FLIM. Nat Commun 5, 3936 (2014).

12. Yang, X., Ha, G. & Needleman, D. J. A coarse-grained NADH redox model enables inference of subcellular metabolic fluxes from fluorescence lifetime imaging. eLife 10, e73808 (2021).

13. Hung, Y. P., Albeck, J. G., Tantama, M. & Yellen, G. Imaging Cytosolic NADH-NAD+ Redox State with a Genetically Encoded Fluorescent Biosensor. Cell Metabolism 14, 545–554 (2011).

14. Zhao, Y. et al. SoNar, a Highly Responsive NAD+/NADH Sensor, Allows High-Throughput Metabolic Screening of Anti-tumor Agents. Cell Metabolism 21, 777–789 (2015).

15. Koberstein, J. N. et al. Monitoring glycolytic dynamics in single cells using a fluorescent biosensor for fructose 1,6-bisphosphate. Proc. Natl. Acad. Sci. U.S.A. 119, e2204407119 (2022).

16. Arce-Molina, R. et al. A highly responsive pyruvate sensor reveals pathway-regulatory role of the mitochondrial pyruvate carrier MPC. eLife 9, e53917 (2020).

17. Li, X. et al. Ultrasensitive sensors reveal the spatiotemporal landscape of lactate metabolism in physiology and disease. Cell Metabolism 35, 200–211.e9 (2023).

18. Nasu, Y. et al. Lactate biosensors for spectrally and spatially multiplexed fluorescence imaging. Nat Commun 14, 6598 (2023).

19. Koveal, D. et al. A high-throughput multiparameter screen for accelerated development and optimization of soluble genetically encoded fluorescent biosensors. Nat Commun 13, 2919 (2022).

20. Wang, Y. et al. Saturation of the mitochondrial NADH shuttles drives aerobic glycolysis in proliferating cells. Molecular Cell 82, 3270–3283.e9 (2022).

21. Kim, W. et al. Polyunsaturated Fatty Acid Desaturation Is a Mechanism for Glycolytic NAD+ Recycling. Cell Metabolism 29, 856–870.e7 (2019).

22. Quinn, W. J. et al. Lactate Limits T Cell Proliferation via the NAD(H) Redox State. Cell Reports 33, 108500 (2020).

23. Luengo, A. et al. Increased demand for NAD+ relative to ATP drives aerobic glycolysis. Molecular Cell 81, 691–707.e6 (2021).

24. Cori, G. T., Slein, M. W. & Cori, C. F. CRYSTALLINE d-GLYCERALDEHYDE-3-PHOSPHATE DEHYDROGENASE FROM RABBIT MUSCLE. Journal of Biological Chemistry 173, 605–618 (1948).

25. Buehner, M., Ford, G. C., Moras, D. & Olsen, K. W. D-Glyceraldehyde-3-Phosphate Dehydrogenase: Three-Dimensional Structure and Evolutionary Significance.

26. Shestov, A. A. et al. Quantitative determinants of aerobic glycolysis identify flux through the enzyme GAPDH as a limiting step. eLife 3, e03342 (2014).

27. Choe, M. & Titov, D. V. Genetically encoded tools for measuring and manipulating metabolism. Nat Chem Biol 18, 451–460 (2022).

28. Pan, X. et al. A genetically encoded tool to increase cellular NADH/NAD+ ratio in living cells. Nat Chem Biol 20, 594–604 (2024).

29. Titov, D. V. et al. Complementation of mitochondrial electron transport chain by manipulation of the NAD^+^ /NADH ratio. Science 352, 231–235 (2016).

30. Goodman, R. P. et al. Hepatic NADH reductive stress underlies common variation in metabolic traits. Nature 583, 122–126 (2020).

31. Yadav, S. et al. Perturbation of NAD(P)H metabolism with the LbNOX xenotopic tool extends lifespan and mitigates age-related changes. Science AdvAnceS (2026).

32. Levine, D. C. et al. NADH inhibition of SIRT1 links energy state to transcription during time-restricted feeding. Nat Metab 3, 1621–1632 (2021).

33. Liu, A. et al. pHmScarlet is a pH-sensitive red fluorescent protein to monitor exocytosis docking and fusion steps. Nat Commun 12, 1413 (2021).

34. Tao, R. et al. Genetically encoded fluorescent sensors reveal dynamic regulation of NADPH metabolism. Nat Methods 14, 720–728 (2017).

35. Hammarlund, M., Hobert, O., Miller, D. M. & Sestan, N. The CeNGEN Project: The Complete Gene Expression Map of an Entire Nervous System. Neuron 99, 430–433 (2018).

36. Jang, S. et al. Glycolytic Enzymes Localize to Synapses under Energy Stress to Support Synaptic Function. Neuron 90, 278–291 (2016).

37. Rangaraju, V., Calloway, N. & Ryan, T. A. Activity-Driven Local ATP Synthesis Is Required for Synaptic Function. Cell 156, 825–835 (2014).

38. Pathak, D. et al. The Role of Mitochondrially Derived ATP in Synaptic Vesicle Recycling. Journal of Biological Chemistry 290, 22325–22336 (2015).

39. Segarra-Mondejar, M. et al. Synaptic activity-induced glycolysis facilitates membrane lipid provision and neurite outgrowth. The EMBO Journal 37, e97368 (2018).

40. Colón-Ramos, D. A., Margeta, M. A. & Shen, K. Glia Promote Local Synaptogenesis Through UNC-6 (Netrin) Signaling in *C. elegans*. Science 318, 103–106 (2007).

41. Liu, S. et al. Glycerol-3-phosphate biosynthesis regenerates cytosolic NAD+ to alleviate mitochondrial disease. Cell Metabolism 33, 1974–1987.e9 (2021).

42. Yao, C.-H. et al. Uncoupled glycerol-3-phosphate shuttle in kidney cancer reveals that cytosolic GPD is essential to support lipid synthesis. Molecular Cell 83, 1340–1349.e7 (2023).

43. Intracellular Oxidation-Reduction States in Vivo. 137, (1962).

44. Williamson, J. R., Scholz, R., Browning, E. T., Thurman, R. G. & Fukami, M. H. Metabolic Effects of Ethanol in Perfused Rat Liver. Journal of Biological Chemistry 244, 5044–5054 (1969).

45. Sharma, R. et al. Circulating markers of NADH-reductive stress correlate with mitochondrial disease severity. Journal of Clinical Investigation 131, e136055 (2021).

46. Thompson Legault, J., et al. A Metabolic Signature of Mitochondrial Dysfunction Revealed through a Monogenic Form of Leigh Syndrome. Cell Reports 13, 981–989 (2015).

47. Singh, C. et al. ChREBP is activated by reductive stress and mediates GCKR-associated metabolic traits. Cell Metabolism 36, 144–158.e7 (2024).

48. Mugabo, Y., et al. Identification of a mammalian glycerol-3-phosphate phosphatase: Role in metabolism and signaling in pancreatic β-cells and hepatocytes.

49. Li, H. et al. Lactate dehydrogenase and glycerol-3-phosphate dehydrogenase cooperatively regulate growth and carbohydrate metabolism during *Drosophila melanogaster* larval development. Development dev.175315 (2019) doi:10.1242/dev.175315.

50. Abdellatif, M., Sedej, S. & Kroemer, G. NAD^+^ Metabolism in Cardiac Health, Aging, and Disease. Circulation 144, 1795–1817 (2021).

51. Zhou, L., Stanley, W. C., Saidel, G. M., Yu, X. & Cabrera, M. E. Regulation of lactate production at the onset of ischaemia is independent of mitochondrial NADH/NAD^+^ : insights from *in silico* studies. The Journal of Physiology 569, 925–937 (2005).

52. Lee, C. F. et al. Normalization of NAD^+^ Redox Balance as a Therapy for Heart Failure. Circulation 134, 883–894 (2016).

53. Thompson Legault, J., et al. A Metabolic Signature of Mitochondrial Dysfunction Revealed through a Monogenic Form of Leigh Syndrome. Cell Reports 13, 981–989 (2015).

54. Zhang, S. et al. Reductive stress: The key pathway in metabolic disorders induced by overnutrition. Journal of Advanced Research 77, 569–584 (2025).

55. Luo, M., Ma, X. & Ye, J. Reductive stress—a common metabolic feature of obesity and cancer. Acta Pharmaceutica Sinica B 14, 5181–5185 (2024).

56. Ross, J. M. et al. High brain lactate is a hallmark of aging and caused by a shift in the lactate dehydrogenase A/B ratio. Proc. Natl. Acad. Sci. U.S.A. 107, 20087–20092 (2010).

57. Zhu, X.-H., Lu, M., Lee, B.-Y., Ugurbil, K. & Chen, W. In vivo NAD assay reveals the intracellular NAD contents and redox state in healthy human brain and their age dependences. Proc. Natl. Acad. Sci. U.S.A. 112, 2876–2881 (2015).

58. Brenner, S. THE GENETICS OF CAENORHABDITIS ELEGANS. Genetics 77, 71–94 (1974).

59. Mello, C. C., Kramer, J. M., Stinchcomb, D. & Ambros, V. Efficient gene transfer in C.elegans: extrachromosomal maintenance and integration of transforming sequences.

60. Paix, A., Folkmann, A., Rasoloson, D. & Seydoux, G. High Efficiency, Homology-Directed Genome Editing in *Caenorhabditis elegans* Using CRISPR-Cas9 Ribonucleoprotein Complexes. Genetics 201, 47–54 (2015).

61. Arribere, J. A. et al. Efficient Marker-Free Recovery of Custom Genetic Modifications with CRISPR/Cas9 in *Caenorhabditis elegans*. Genetics 198, 837–846 (2014).

62. Crane, M. M. et al. In vivo measurements reveal a single 5′-intron is sufficient to increase protein expression level in Caenorhabditis elegans. Sci Rep 9, 9192 (2019).

