## Supplementary Tables for "A Multisensor Framework Reveals Redox Constraints on Glycolysis *in vivo*"

### **SUPPLEMENTAL TEXT**

**Supplementary Table S1: STRAIN LIST**

| **Number** | **Strain name** | **Genotype** | **Strain description** |
| --- | --- | --- | --- |
| 1 | DCR9694 | *Ex5680* | Pttx-3::FiLa (FiLa expressed in AIY) |
| 2 | DCR10038 | *Ex5680; pfk-1.1 (ola458)* | Pttx-3::FiLa; pfk-1.1(-) |
| 3 | DCR9695 | *Ex5680; ldh-1(ola514)* | Pttx-3::FiLa; *ldh-1(-)* background |
| 4 | DCR10005 | *olaEx5725* | Pttx-3::FiLaC (Reduced affinity variant FiLa-C expressed in AIY) |
| 5 | DCR10016 | *olaEx5728* | Pttx-3::pHmScarlet |
| 6 | DCR10039 | *olaEx5728; ldh-1(ola514)* | pHmScarlet in an ldh-1(-) background |
| 7 | DCR9692 | *Ex5678* | SoNar expressed in AIY |
| 8 | DCR10006 | *olaEx5726* | Reduced affinity variant iNapc expressed in AIY |
| 9 | DCR10040 | *Ex5678; Ex5679* | Pttx-3::SoNar; Pttx-3::EcSTH |
| 10 | DCR9728 | *Ex5678; Ex5685* | Pttx-3::SoNar; Pttx-3::LbNOX |
| 11 | DCR10017 | *Ex5678; ldh-1 (ola514)* | Pttx-3::SoNar; ldh-1(-) |
| 12 | DCR10018 | *Ex5678; mdh-1(ola599)* | Pttx-3::SoNar; mdh-1(-) |
| 13 | DCR10047 | *Ex5678; gpdh-2(ola518)* | Pttx-3::SoNar; gpdh-2(-) |
| 14 | DCR10048 | *Ex5680; mdh-1(ola599)* | Pttx-3::FiLa; mdh-1(-) |
| 15 | DCR9726 | *Ex5680; gpdh-2(ola518)* | Pttx-3::FiLa; gpdh-2(-) |
| 16 | DCR9696 | *olaIs138; Ex5679* | Pttx-3::HYlight; Pttx-3::EcSTH |
| 17 | DCR10049 | *olaIs138; Ex5685* | Pttx-3::HYlight; Pttx-3::LbNOX |
| 18 | DCR9727 | *Ex5680; Ex5679* | Pttx-3::FiLaR; Pttx-3::EcSTH |
| 19 | DCR10050 | *Ex5680; Ex5685* | Pttx-3::FiLaR; Pttx-3::LbNOX |
| 20 | DCR9133 | *olaex5431* | Pttx-3::Pyronic-SF |
| 21 | DCR9134 | *olaex5432* | Pttx-3::Pyronic-SFdead |
| 22 | DCR10052 | *olaex5431; Ex5685* | Pttx-3::Pyronic-SF; LbNOX |
| 23 | DCR10053 | *olaex5432; Ex5685* | Pttx-3::Pyronic-SFdead; LbNOX |
| 24 | DCR8818 | *olaIs123* | Pttx-3::RAB-3::Mcherry |
| 25 | DCR9715 | *olaIs123; ldh-1(ola514)* | Pttx-3::RAB-3::Mcherry; ldh-1(-) |
| 26 | DCR9693 | *Ex5679; olaIs123* | Pttx-3::RAB-3::Mcherry; Pttx-3::EcSTH |
| 27 | DCR10054 | *olaIs123; gpd-2 gpd-3(ola532)* | Pttx-3::RAB-3::Mch; gpd-2(-) gpd-3(-) |
| 28 | DCR9718 | *olaIs138; gpd-2 gpd-3(ola532)* | Pttx-3::HYlight; gpd-2(-) gpd-3(-) |
| 29 | DCR9712 | *olaIs138; ldh-1(ola514)* | Pttx-3::HYlight; ldh-1(-) |
| 30 | DCR9713 | *olaIs138; gpdh-2(ola518)* | Pttx-3::HYlight; gpdh-2(-) |
| 31 | DCR10013 | *olaIs138; mdh-1(ola599)* | Pttx-3::HYlight; mdh-1(-) |
| 32 | DCR9714 | *olaIs138; gpdh-2(ola518); ldh-1 (ola514)* | Pttx-3::HYlight; gpdh-2(-); ldh-1(-) |
| 33 | DCR10057 | *Ex5678; gpdh-2(ola518); ldh-1 (ola514)* | Pttx-3::SoNar; gpdh-2(-); ldh-1(-) |
| 34 | DCR10064 | *Ex5678; ldh-1(ola514); Ex5681* | Pttx-3::SoNarR; ldh-1(-); Pttx-3::GPDH-2BcDNA |
| 35 | DCR9700 | *olaIs138; olaEx5681; ldh-1(ola514)* | Pttx-3::HYlight; ldh-1(-) ; Pttx-3::GPDH-2B cDNA |
| 36 | DCR9704 | *ldh-1(ola514); gpd-2 gpd-3 (ola532); olaEx5681; olaIs138* | Pttx-3::HYlight; ldh-1(-); gpd-2(-) gpd-3(-); Pttx-3::GPDH-2B cDNA |
| 37 | DCR10056 | *olaIs123; Ex5685* | Pttx-3::RAB-3::Mcherry; LbNOX |
| 38 | DCR9716 | *olaIs123; gpdh-2(ola518)* | Pttx-3::RAB-3::Mcherry; gpdh-2(-) |
| 39 | DCR10055 | *olaIs123; mdh-1(ola599)* | Pttx-3::RAB-3::Mcherry; mdh-1(-) |
| 40 | DCR9910 | *olaEx5705* | ASE::R-iLACCO1.2. |
| 41 | DCR9935 | *ldh-1(ola514); olaEx5705* | ASE::R-iLACCO1.2; ldh-1(-) |
| 42 | DCR9933 | *olaEx5710* | ASE:: FiLa::mScarlet-I3 |
| 43 | DCR9934 | *ldh-1(ola514); olaEx5710* | ASE:: FiLa::mScarlet-I3; ldh-1(-) |
| 44 | DCR9089 | *olaIs138* | HYlight expressed in AIY |
| 45 | DCR9697 | *ldh-1(ola514); olaEx5681; olaIs123* | Pttx-3::RAB-3::Mcherry; ldh-1(-); Pttx-3::GPDH-2B cDNA |

**Supplementary Table S2:PLASMID LIST**

| **Number** | **Plasmid Name** | **Description** |
| --- | --- | --- |
| 1 | DACR4153 | Pttx-3::SoNar::T2A::Mscarlet-I3::unc-54 3’TUR |
| 2 | DACR4186 | Pttx-3::FiLa::T2A::Mscarlet-I3::unc-54 3’TUR |
| 3 | DACR4187 | Pttx-3::iNapc::unc-54 3’TUR |
| 4 | DACR4188 | Pttx-3::FiLaC::unc-54 3’TUR |
| 5 | DACR4189 | Pttx-3::pH sensor::unc-54 3’TUR |
| 6 | DACR4190 | Pttx-3::Pyronic-SF::unc-54 3’TUR |
| 7 | DACR4191 | Pttx-3::Pyronic-SF-NA::unc-54 3’TUR |
| 8 | DACR4149 | Pttx-3::EcSTH::unc-54 3’TUR |
| 9 | DACR4151 | Pttx-3::LbNOX::unc-54 3’TUR |
| 10 | DACR4192 | Pttx-3::GPDH-2B::unc-54 3’TUR |
| 11 | DACR4193 | Pfl-6::FiLa::T2A::MscarletI3::let-858 3’TUR |
| 12 | DACR4194 | Pflp-6::RiLACCO::T2A::BFP::unc-10 3’ |

**Supplementary Table S3: OLIGONUCLEOTIDE LIST**

| **Number** | **Oligo sequence** | **Description** |
| --- | --- | --- |
| P01 | CGCGCATTCATTGTCCGCATC | Triplex primers to genotype *pfk-1.1 (ola458)* |
| P02 | GATCCATCTCCACCAATGCATACAAG | Triplex primers to genotype *pfk-1.1 (ola458)* |
| P03 | GGCTAAGACTTCAATCAACTTGTGCAAA | Triplex primers to genotype *pfk-1.1 (ola458)* |
| P04 | TCGAAAGCTCACCACATCATC | Triplex primers to genotype *ldh-1 (ola514)* |
| P05 | GGCGAAAGTGAGAGATAGAAG | Triplex primers to genotype *ldh-1 (ola514)* |
| P06 | GACAAAGCGGTAGGGAATAG | Triplex primers to genotype *ldh-1 (ola514)* |
| P07 | CAACGACTGAAGGAAATCGAG | Triplex primers to genotype *gpd-2 /gpd-3(ola532)* |
| P08 | AAGACGAGCAGTGAGATCAAC | Triplex primers to genotype *gpd-2 /gpd-3(ola532)* |
| P09 | TGACGGACAATCACAACGAAG | Triplex primers to genotype *gpd-2 /gpd-3(ola532)* |
| P10 | CATCTGCGTCTCCACTATTC | Triplex primers to genotype *gpdh-2 (ola518)* |
| P11 | AATTGCATAGGGAGTCCAAGC | Triplex primers to genotype *gpdh-2 (ola518)* |
| P12 | GTGTTCTGGATGGTTTCTGAG | Triplex primers to genotype *gpdh-2 (ola518)* |
| P13 | CCGTAGGGTTTGTTGACAAG | Triplex primers to genotype *mdh-1 (ola599)* |
| P14 | TGATCTCACCTTGGTGGTTG | Triplex primers to genotype *mdh-1 (ola599)* |
| P15 | TGTTAATCTGAGTGGGTCTCG | Triplex primers to genotype *mdh-1 (ola599)* |
| P16 | TGCATAATTCGAAATGGACG | crRNA to generate *pfk-1.1(ola458)* |
| P17 | TTTTCAGAACATGTGTGAGT | crRNA to generate *pfk-1.1(ola458)* |
| P18 | ATTGTCATAAATTACAGAATTTTGCATAATTCGAACACACATGTTCTGAAAAGTTTTGATGTCTCCATAG | Homology template to generate *pfk-1.1(ola458)* |
| P19 | GATATCTATTAAAAATGACA | crRNA to generate *ldh-1(ola514)* |
| P20 | GTTCGATGACTGAAGGAAAA | crRNA to generate *ldh-1(ola514)* |
| P21 | AAATACTCTACTGCAATTGCTGATATCTATTAAAAAACTGTTTATATAGCATACTTCCCCTTGTCACAGA | Homology template to generate *ldh-1(ola514)* |
| P22 | TTTCTGACATCAGATTTTCT | crRNA to generate *gpdh-2(ola518)* |
| P23 | GATGATGCACAAGACTGGAT | crRNA to generate *gpdh-2(ola518)* |
| P24 | TTTTGTTTTCAGAAAAAGTTTCTGACATCAGATTTGATTGGATGCTAAGTGAGTACTAGAAATTATTATT | Homology template to generate *gpdh-2(ola518)* |
| P25 | GACCAAAACGCGAAGTGGGG | crRNA to generate *mdh-1(ola599)* |
| P26 | TATGGGGAAAATTTAGATGT | crRNA to generate *mdh-1(ola599)* |
| P27 | ACACATTTCTTAGTGTAAATTGCTAAATTTTCAGATAAATTTTCCCCATAAATTATAACGATTTCCCAAC | Homology template to generate *mdh-1(ola599)* |
| P28 | TCCGACACTTGGCTTTGGCA | crRNA to generate *gpd-2/gpd-3(ola532)* |
| P29 | TAGAGAATTTGAAGCGGCTT | crRNA to generate *gpd-2/gpd-3(ola532)* |
| P30 | AGTTGAACTGATTGTAGTAGTATCAATTTTCAGCCGCCGCTTCAAATTCTCTAAGTCTGAAATGAATCTG | Homology template to generate *gpd-2/gpd-3(ola532)* |
